# Dynamic Flux Balance Analysis Models in SBML

**DOI:** 10.1101/245076

**Authors:** Matthias König, Leandro H. Watanabe, Jan Grzegorzewski, Chris J. Myers

## Abstract

Computational models in systems biology and systems medicine are typically simulated using a single formalism such as ordinary differential equations (ODE). However, more complex models require the coupling of multiple formalisms since different biological phenomena are better described by different methods. For example, metabolism in steady state is often modeled using flux-balance analysis (FBA) whereas dynamic changes of model components are better described via ODEs. The coupling of FBA and ODE modeling formalisms results in dynamic FBA models. A major challenge is how to describe such hybrid models that couple multiple formalisms in a standardized way so that they can be exchanged between tools and simulated consistently in a reproducible manner. This paper presents a scheme for encoding and implementation of dynamic FBA models in the Systems Biology Markup Language (SBML), thereby enabling the exchange of multi-framework computational models between software tools. We demonstrate the feasibility of the approach using various example models and show that different tools are able to simulate the hybrid models and agree on the results. As part of this work, two independent implementations of a multi-framework simulation method for dynamic FBA have been developed supporting such models: `iBioSim` and `sbmiutils`.

## 1 INTRODUCTION

In systems biology, mathematical modeling is used to investigate biological systems (Kitano, 2002). The resulting computational models enable researchers to make predictions *in silico* which can be validated experimentally. However, the process of model building is time-consuming and error-prone. Model reproducibility and exchangeability are of major importance for independent validation of results and model reuse, especially in the case of more complex models.

To achieve reproducibility, interoperability, and consistent model interpretation, a well-defined modeling representation with unambiguous syntax is crucial. To this end, standard model representation formats exist that enable model exchange, such as the Systems Biology Markup Language (SBML) (Hucka et al., 2003; Keating et al., 2020; Hucka et al., 2019).

SBML has been successfully applied to the encoding of single formalism models, but the encoding of hybrid models using SBML has yet to be explored. Some tools have implemented hybrid simulation, such as COPASI (Hoops et al., 2006) and E-CELL (Tomita et al., 1999), nonetheless, they lack reproducibility. In COPASI, the models fall short of necessary pieces of information for model exchange. Namely, these models lack the information about their own model formalism which results in hybrid models being only specific to COPASI. In E-CELL, most models are encoded in C++ and only few in SBML. Even though the C++ models are repeatable, they are not reproducible because other tools cannot use these files and even models encoded in SBML are incomplete and lack the integration of different formalisms.

The support of hybrid modeling adds new challenges. The present work addresses this problem by developing a methodology in conjunction with implementations to support such hybrid modeling efforts. We demonstrate the usefulness of our approach by exchanging two models between two distinct simulation tools with both implementations leading to similar simulation results.

### 1.1 Coupling multiple modeling formalisms

Various simulation and analysis methods have been developed in systems biology. Depending on the biological question, different methods are preferred. Kinetic time-course simulations based on ordinary differential equations (ODE) are often employed to study the dynamics of entities in a model over time. Depending on the research question and biological system, such simulations can be non-deterministic (stochastic). Other popular simulation methods are Boolean (Thomas, 1973; Kauffman, 1969) models, logical models (Morris et al., 2010), and constraint-based approaches (Bordbar et al., 2014).

Dynamical modeling of metabolic networks by ODE approaches is particularly challenging since kinetic parameters needed for ODE models are often unobtainable (Varma and Palsson, 1994). Hence, steady-state approaches that do not need kinetic information are employed to model metabolism such as flux balance analysis (FBA) (Savinell and Palsson, 1992; Varma et al., 1993) which is based on constraint-based optimization. This method only requires the connectivity of the reactions and metabolites along with the respective stoichiometry, an objective function, such as cell growth, and additional constraints like flux bounds. The idea is to constrain the model based on the stoichiometry of the reactions and optimize the objective function while satisfying the flux constraints. These models do not require kinetic information and can be simulated efficiently even in case of very large systems.

Biological research questions often require the coupling of different model formalisms. One such recent example is the whole-cell model for the *Mycoplasma genitalium* (Karr et al., 2012) that is encoded using a mixture of Boolean networks, stochastic processes, differential equations, and FBA.

### 1.2 Dynamic flux balance analysis

One disadvantage of FBA is that it cannot express the dynamics of the metabolites since it does not change amounts or concentrations of species, but only provides information about the optimal flux distribution for the given optimization problem. Due to this limitation, the field of dynamic FBA (DFBA) (Varma and Palsson, 1994) has emerged, which couples the stationary flux distribution resulting from FBA with the kinetic update of the metabolites taken up or consumed by the FBA network. For DFBA models, a FBA submodel is coupled to a kinetic model (ODE).

Besides the whole-cell model, which uses DFBA as a core module, various metabolic pathways have been modeled using DFBA. DFBA has been applied in small-scale examples (Varma and Palsson, 1994; Mahadevan et al., 2002; Luo et al., 2006), over medium-size models (Pizarro et al., 2007; Lequeux et al., 2010; Meadows et al., 2010), and up to genome-scale DFBA applications (Hanly and Henson, 2011; Hjersted et al., 2007). For an overview, see Table 1 in (Höffner et al., 2013).

The coupling between FBA and kinetic model parts can be implemented via three main approaches: static optimization approach (SOA), dynamic optimization approach (DOA), and direct approach (DA) (Gomez et al., 2014). The SOA approach solves the linear programming (LP) problem of FBA at each time step using an Euler forward method assuming constant fluxes over the time step (Gomez et al., 2014). DOA approaches optimize simultaneously over the entire time period by solving a nonlinear programming problem (NLP). The DA approach directly includes the LP solver on the right-hand side of the ordinary differential equations (ODEs).

The advantage of the SOA is its relatively simple implementation, which is why most of the published DFBA models use the SOA approach. However, SOA is often less accurate compared to other computational more demanding methods such as DOA. The DA method exhibits the best trade-off between accuracy and runtime performance but has its downsides in terms of implementation difficulty. For this work, we use the SOA method. Its simplicity makes it a good candidate to use as proof of concept for this work.

### 1.3 Exchangeability & reproducibility of models

Despite the wide range of published DFBA models no standard for the exchange of such models exists. Existing models are hard-coded, such as the whole-cell model which is implemented in MATLAB. Hereby, the mathematical model is separated in the respective kinetic and FBA formalisms in a script along with the connections between the kinetic and flux balance parts of the models. As a consequence, it is not possible to exchange existing DFBA models between different software tools. Thus, they cannot be reproduced or validated. This is especially problematic in the case of DFBA models because often multiple optima can exist for the FBA model part (and the various time steps). The resulting DFBA results are not unique since they depend on the analysis implementation (how a solver selects one of the possible FBA solutions). In addition, the simulation results may depend on the selected step size of the SOA algorithm, in particular, if the step size is not small enough.

While it is possible to replicate the same scripts in different programming languages, it is unpractical, error-prone, unnecessary, often leads to data loss, and most importantly does not solve the underlying problem of non-exchangeability of such models. For these reasons, script replication makes achieving reproducibility difficult and often infeasible. The necessity of an exchange format for DFBA emerged from efforts trying to encode and reproduce the DFBA submodel of the whole-cell model using standards during the whole-cell workshop (Waltemath et al., 2016).

### 1.4 Model standards

To achieve exchangeability and reproducibility of models, standards for the encoding of models have been created. The de-facto standard for systems biology models is SBML (Hucka et al., 2003; Keating et al., 2020). SBML core elements are used to describe mathematical models of reaction-based networks and provide the means to encode computational models based on reaction networks that can be represented both deterministically and stochastically. SBML uses packages for extending the functionality of core elements. While SBML is used to encode mathematical models of biological networks, there are different standards for other purposes: the Simulation Experiment Description Markup Language (SED-ML) is used for describing simulations (Waltemath et al., 2016; Bergmann et al., 2018), the Systems Biology Graphical Notation (SBGN) is used for describing visualizations (Le Novère et al., 2009), and COMBINE Archives are used for grouping files in a single archive necessary to reproduce a modeling experiment (Bergmann et al., 2014). The main advantage of using these standards over hard-coding models in code is the ability to exchange models between research groups and reproduce results using various tools that support these standards. In addition, these standards enable the use of semantic annotations to document the model and model components which enhances the reusability and interoperability (Neal et al., 2019, 2020).

One of the challenges in SBML models is the limitation of models to a single formalism lacking support for the expression of models using multiple formalisms. Although there are several tools that support ODE simulation and FBA, they all support them independently. In order to overcome this challenge, this paper introduces a scheme that enables the coupling of ODE and FBA models. This paper demonstrates that this scheme facilitates exchangeability and reproducibility by encoding and simulating DFBA models in both `iBioSim` (Watanabe et al., 2019) and `sbmlutils` (König, 2022).

## 2 MATERIAL AND METHODS

### 2.1 Model encoding

The DFBA models presented in this paper were created in the proposed scheme either using a graphical user interface in `iBioSim` or a script-based approach in `sbmlutils`. For a given model, the four submodels (TOP, FBA, BOUNDS, and UPDATE) were packaged with the corresponding simulation files using SED-ML in COMBINE archives in order to facilitate the exchange between tools. All models and simulation results are available from https://github.com/matthiaskoenig/dfba.

### 2.2 Stationary optimization approach (SOA)

A stationary optimization approach for DFBA was implemented as a simulation algorithm in `iBioSim` and `sbmlutils` following the simulation scheme depicted in Figure 1.

**Figure 1.**
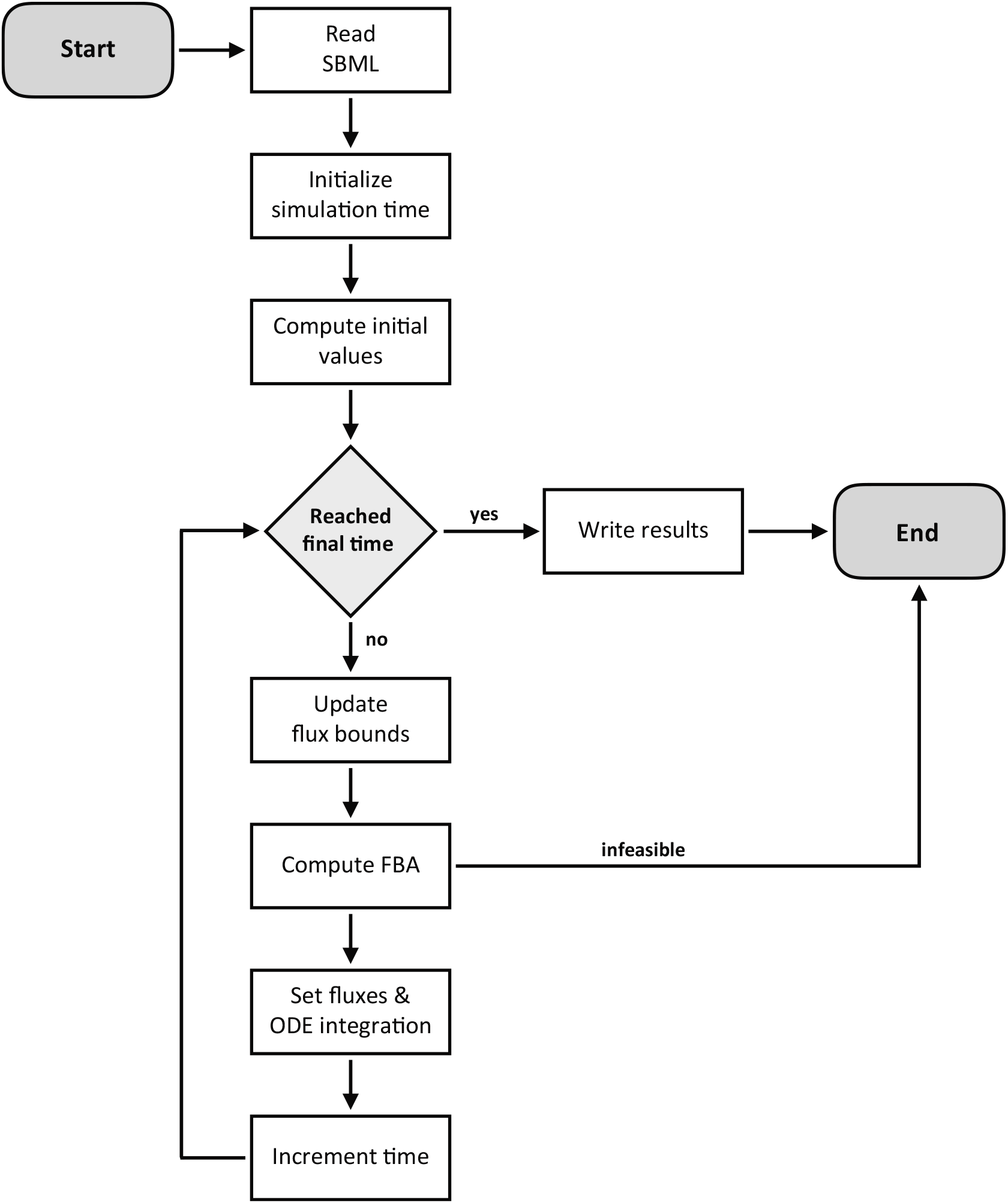
Overview of the implemented SOA algorithm for DFBA. After the initialization of the model, the FBA and kinetic simulations are run iteratively until the simulation endpoint. In every step, FBA is used to compute the reaction rates of the FBA network. Subsequently, based on the computed FBA rates, the values of the species are updated dynamically. In the SOA approach, FBA fluxes are assumed to be constant within a time step. For a detailed description see the Material And Methods Section.

The following paragraph assumes familiarity with SBML and we refer to the SBML specification for more information (Hucka et al., 2019). As the first step, all of the species and parameters in the model are initialized and each variable is assigned an initial value. After the initialization step, the FBA submodel is executed. During the FBA step, reaction fluxes are computed using the initial flux bound values where the flux bounds for the reactions come from the top-level using replacements from the SBML comp package (Smith et al., 2015). In SBML, replacements of parameters and species indicate the top-level entities are the same entity as the one being replaced. Once the fluxes are computed, they are assigned on the top-level to parameters using assignment rules. These parameters represent reaction rates.

After computing reaction fluxes, the update step is performed concurrently with a dynamic step by computing the time-evolution of every species in the UPDATE and KINETIC submodels. Species that affect any flux bound in the FBA submodel are updated on the top-level. The new bounds are used in the FBA submodel for the next time step. Simulation time is incremented at the end. If the time limit is reached, then the simulation is complete. Otherwise, all of the steps are repeated.

The SOA simulation algorithm has been implemented in `iBioSim` and `sbmlutils`. The `iBioSim` tool uses the structure of (Watanabe and Myers, 2014) for simulation. The `sbmlutils` tool uses roadrunner (Somogyi et al., 2015) for the kinetic simulation and cobrapy (Ebrahim et al., 2013) to solve the FBA problem. Both `iBioSim` and `sbmlutils` take an SBML file that describes a DFBA model and a SED-ML file that describes the simulation experiment. In the proposed approach, SED-ML is mainly used to indicate which simulation algorithm to use, the time points in which tools should print out the values of the variables, the initial time, and the time limit. The SED-ML files provide a minimal simulation experiment to check reproducibility between implementations. The value of each time increment for SOA is defined by a parameter with id `dt` in the SBML model, which can be overwritten by the SED-ML file for the actual simulation. An ontology term for the description of DFBA simulation algorithms has been introduced in the Kinetic Simulation Algorithm Ontology (KISAO) (Zhukova et al., 2011), term `KISAO:0000500` corresponding to the DFBA-SOA method, and is used in the SED-ML descriptions.

### 2.3 Reproducibility between tools

In order to test interoperability based on the proposed scheme, models were built in both the `iBioSim` and the `sbmlutils` tools. Models built in `iBioSim` were then imported into `sbmlutils` and vice-versa to check whether models could be interpreted by both tools consistently. This was done in an iterative manner and issues were solved by clarifying the encoding scheme by adding additional rules which resolved ambiguities. Ensuring reproducibility for DFBA models is challenging because there may exist several possible outcomes that satisfy the objective function and constraints of the FBA models. Different trajectories can result from the DFBA simulation depending on how a solver and implementation selects one of the multiple optima. The issue of multiple optima was solved by guaranteeing uniqueness of the solution in every time step based on Flux Variability Analysis (FVA) (Mahadevan and Schilling, 2003). FVA gives the possible minimal and maximal fluxes for each reaction in each step of the simulation. If all minimal fluxes are equal to all maximal fluxes for a time point a solution is unique in the time point. If all time points are unique the solution is unique. As a practical note: If the solution is not unique, the addition of additional constraints to the FBA problem allows to make the solution unique. Reproducibility of the model simulations was tested by comparing the numerical SOA results between the two tools for models with unique solutions (see Supplementary Material S2). Results were assumed as numerically identical if the absolute difference for every time point *t_k_* for all dynamical FBA species in the model *c_k_* was smaller than the tolerance *ϵ* = 10^-5^. The difference is computed as follows:

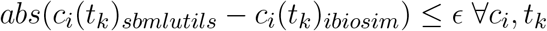

In SOA-DFBA, it is important that the time steps `dt` are small enough so that the solution converges against the optimal solution. Solutions vary if selected step sizes are too large. To highlight this fact, changing the step size in the `toy_wholecell` model from 1.0 to 0.1 resulted in differences in steady-state concentrations of up to 10%. Consequently, the step size was reduced until the changes did not affect the simulation results.

## 3 RESULTS

The major result of this work is the creation of the first schema for a DFBA encoding in SBML, demonstrating hybrid computational models to be exchangeable and reproducible between tools. In the following, the schema and its application to multiple DFBA models is presented.

### 3.1 Schema for dynamic flux balance analysis

This paper proposes for the first time a schema to encode hybrid models, such as DFBA model, in SBML. The developed schema consists of rules, guidelines, and additional information and is available in the Supplementary Material S1. The latest version of the document is available from https://github.com/matthiaskoenig/dfba/. Proposals, errata, and updates to the schema are managed via the respective issue tracker and releases.

In this Section, we provide a high-level overview of the underlying concepts used in the schema, followed by an application of the schema to encode DFBA models.

The DFBA model is constructed hierarchically using the SBML comp package, separating the hybrid model into different building blocks based on the respective functionality and modeling frameworks (Figure 2). The top-level model is hereby composed of four submodels: (i) a kinetic submodel that computes flux bounds based on the dynamic metabolite availability and ensures that the FBA problem is constrained by the available metabolite resources (BOUNDS submodel); (ii) a FBA submodel that encodes metabolism as a FBA problem (FBA submodel); and (iii) a kinetic submodel that updates the amounts and concentrations of the dynamic metabolites changed via the FBA submodel via consumption or production (UPDATE submodel); (iv) an optional kinetic submodel that represents a dynamic part with all kinetics other than the metabolic pathway, such as DNA transcription, DNA translation, and protein degradation, among others (KINETIC submodel). Alternatively, arbitrary kinetics can be part of the top model.

**Figure 2.**
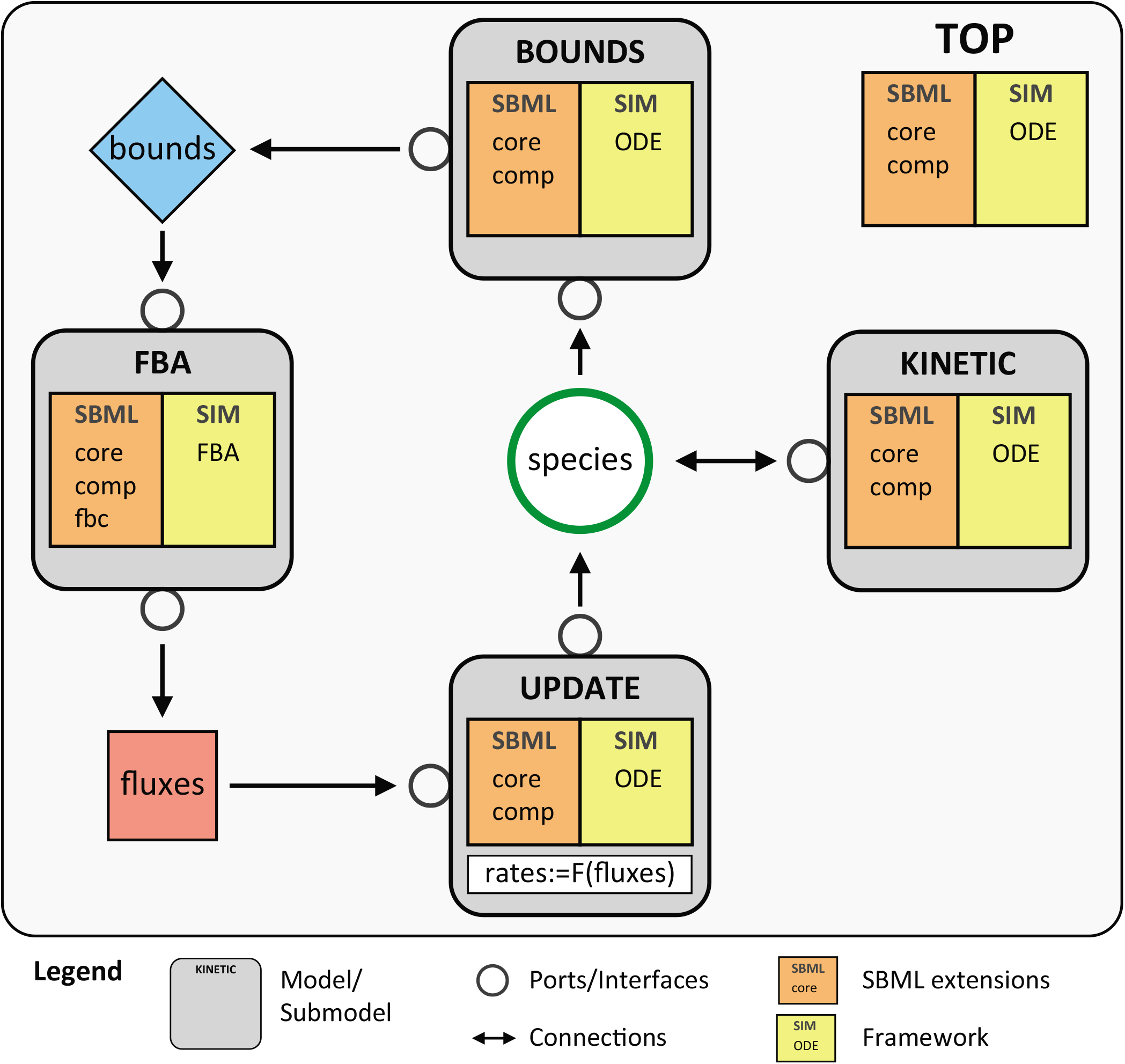
Overview of the schema for DFBA model encoding in SBML. The hierarchical SBML model is composed of a top-level model with four submodels: FBA, BOUNDS, UPDATE, and KINETIC. The individual submodels are connected via ports. The respective SBML packages used are listed in the models, as well as the employed simulation method. The BOUNDS submodel calculates the upper and lower flux bounds based on metabolite availability. The FBA submodel computes the reaction fluxes of the metabolic model encoded via the SBML fbc package using the bounds as constraints. The UPDATE submodel calculates the dynamic update of the dynamic metabolites affected by the FBA model. The rates of change are hereby functions of the FBA fluxes. The KINETIC submodel includes all of the other processes in the model, which may affect or be affected by entities in metabolism. The top-level model ties together the different submodels using SBML comp `replacedElement` and `replacedBy` constructs.

**Figure 3.**
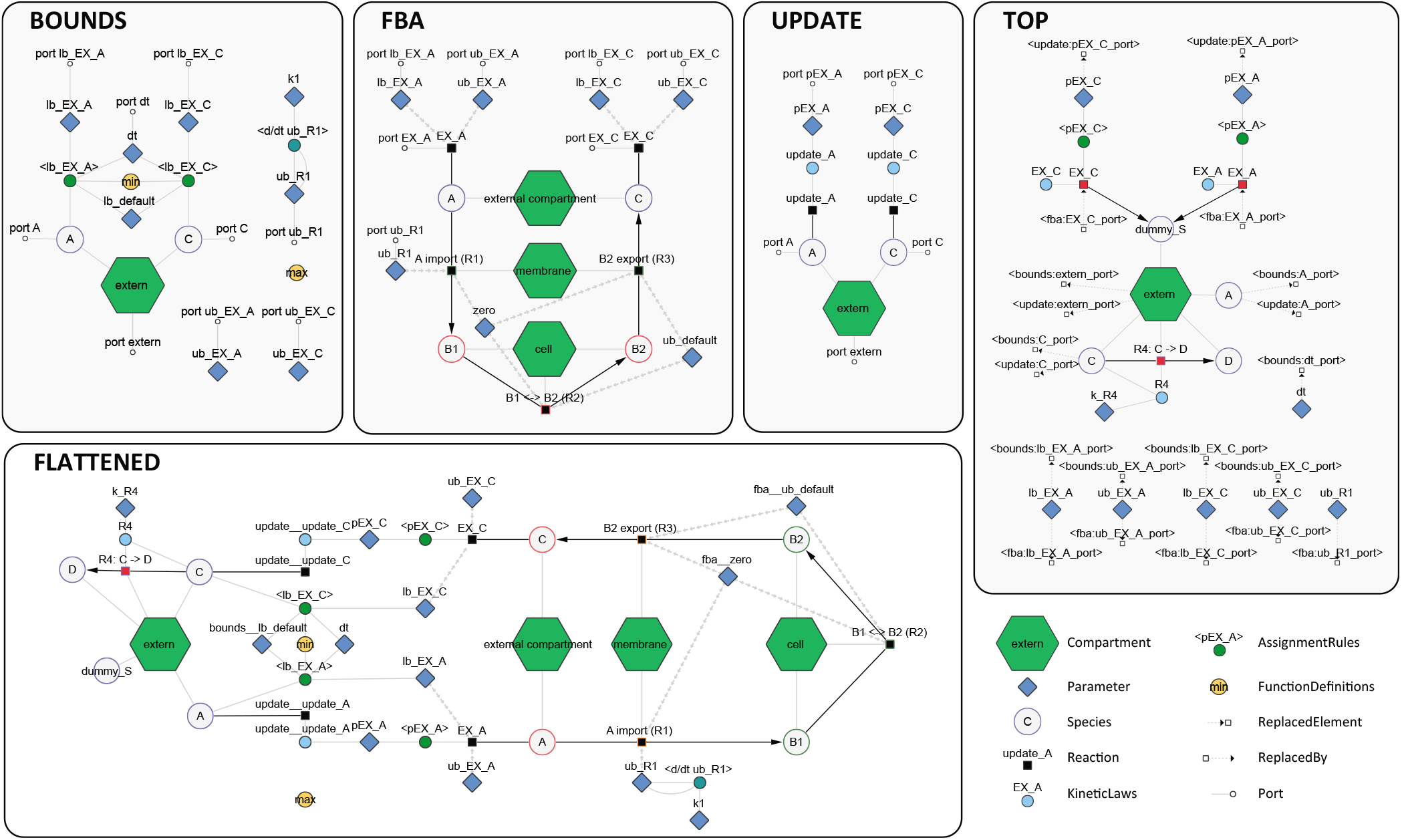
Detailed schema of the minimal example model (`toy_wholecell`). The figure shows the components in the BOUNDS, FBA and UPDATE submodels. Links between submodel components are based on ports which are connected elements via TOP model replacements (`replacedElements` and `replacedBy`). The flattened SBML comp model (FLATTENED) shows the resolved connections between the different submodels after these replacements have been performed. The flattened model can not be simulated because the separation of the modeling formalisms is lost in the flattening process. The network visualization are available as interactive graphs in Cytoscape as Supplementary Material, which provide additional information and annotation of the components. The figure was created with cy3sbml using the SBML models (König et al., 2012).

The top-level model couples the three different submodels using SBML comp `replacedElement` and `replacedBy` constructs with the interface between the submodels defined via comp ports (which define which model components of the submodels can be connected, i.e, are exposed).

In order to describe the different formalisms of each submodel, the Systems Biology Ontology (SBO) is used (Courtot et al., 2011). The SBO defines controlled vocabulary terms used in the systems biology field. The SBO terms are arranged in a taxonomic hierarchy using a tree structure. This allows the grouping of terms that are related to one another. The modeling formalisms of the individual submodels are described using terms on the modeling framework branch, where FBA models are described using the flux balance framework term, stochastic processes are described using the non-spatial discrete framework term, and differential equations are described using the non-spatial continuous framework term. Although the terms for stochastic processes and differential equations can be used for describing either stochastic or deterministic simulation methods, these terms were selected because they are the ones that best describe these two formalisms.

In addition to the modeling formalism, other key components are annotated in the submodels via SBO terms in the schema, like the upper and lower flux bounds and the exchange reactions in the FBA submodel defining which metabolites can be consumed or produced in the FBA part of the DFBA, or the dynamic species in the top model changed by the FBA submodel. By the means of these annotations, the interface between the hybrid submodels can be clearly defined.

All of the interconnections between the submodels are encoded in SBML rather than using an external approach like for instance via SED-ML. The connections between model components are crucial information of the model and should be part of the model encoding. SED-ML is only used to encode which simulation to run with the model. As a consequence, this schema requires only a single hierarchical SBML model and a single SED-ML file.

### 3.2 Minimal Example (toy wholecell)

In order to illustrate the proposed schema, a simplified example of a whole-cell model was created and visualized. The corresponding files (i.e. COMBINE archive and Cytoscape visualization) are in Supplementary Material S3. The visualization shows how the different submodels connect with each other in a flat form.

This model is constructed hierarchically where a top-level model is created to instantiate different submodels (BOUNDS, UPDATE, and FBA) and make the necessary connections between them. The figure illustrates the structure of each submodel and how each submodel ties in with each other in a flat version of the model once all of the connections are established.

In the example, the FBA submodel imports species `A` and convert it via a linear chain of reactions to species `C`. The exchange reactions `EX_A` and `EX_C` contain the rate of consumption and production of the respective species. The TOP model contains assignment rules that assign the fluxes to the parameters `pEX_A` and `pEX_C`. The `pEX_A` and `pEX_C` parameters are used by the UPDATE model to compute the new values of the dynamic species `A` and `C` via the update reactions `update_A` and `update_C`. The BOUNDS model calculates the bounds of all FBA exchange reactions (constraining the availability of the dynamic species). In the example, the upper bound `ub_R1` of reaction `R1` is changed via a rate rule. Additional kinetics are encoded in the TOP model, such as the kinetic conversion of `C` to `D` (these could also be in a separate KINETIC submodel).

In order to validate the exchangeability and reproducibility of the model, simulations were performed using the simulation algorithm described in Figure 1 with results depicted in Figure 4. Both implementations resulted in numerically identical results (see Section 2.3). Importantly, our encoding schema allowed to reproduce the numerical results even if the step sizes were not yet small enough to have converged against the correct solution, thereby allowing to test the effects of varying step sizes in a reproducible manner.

**Figure 4.**
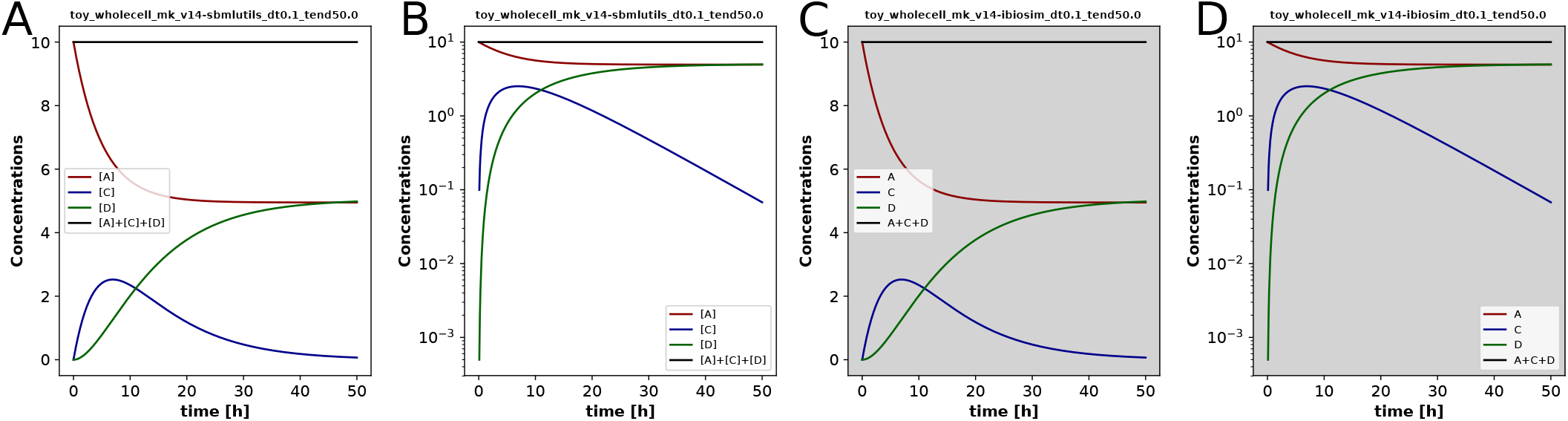
DFBA simulation results for the `toy_wholecell` model in two different tools (`sbmlutils`: **A, B**; `iBioSim`: **C, D**). This demonstrates that models can be exchanged by different tools using standards and the results can be reproduced when using the same simulation algorithm. Species `A` is converted to `C` via the FBA subnetwork over time. Species `C` is converted to `D` via the kinetic parts in the top model. Species `A` is not consumed completely because of the import of `A` in the FBA subnetwork via `R1` which is shut down over time via a rate rule for the upper flux bound. The model was simulated for 50[h] with a time step `dt` of 0.1[h].

In addition to the presented minimal model, a second model and its corresponding Cytoscape visualization of a simplified DFBA glycolysis (`toy_atp`) is available in the supplement (COMBINE archive in Supplementary Material S3)

### 3.3 Diauxic growth in *Escherichia coli* (diauxic growth)

The next example is an encoding and reproduction of results from a published DFBA model of diauxic growth of the *Escherichia coli* (Mahadevan et al., 2002) consisting of four reactions between four metabolites: glucose (*Glcxt* oxygen (*O*_2_), acetate (*A_c_*), and biomass (*X*). The model can grow either aerobically on acetate (*v*1), aerobically on glucose (*v*2 or *v*3), or anaerobically convert glucose to acetate:

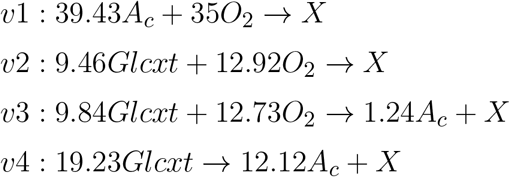

The kinetic part of the model is described by the following differential equations:

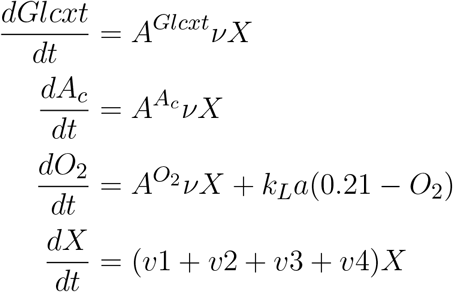

where *A^Glcxt^, A^A_c_^, A*^*O*_2_^ are the respective rows of each variable in the stoichiometry matrix and *k_L_a* is the mass transfer coefficient of oxygen. For a detailed description see (Mahadevan et al., 2002).

The model and its corresponding Cytoscape visualization is available in Supplementary Material S3.

The results in Figure 5 depict an exponential growth phase using glucose aerobically until running out of glucose, which at this point the cell grows linearly due to oxygen. When both oxygen and glucose run out, the cell growth stagnates. Experimental data from (Varma and Palsson, 1994) is plotted alongside the simulation results. The model is able to capture the behavior observed in the experimental data. The results are equivalent to the models in (Mahadevan et al., 2002).

**Figure 5.**
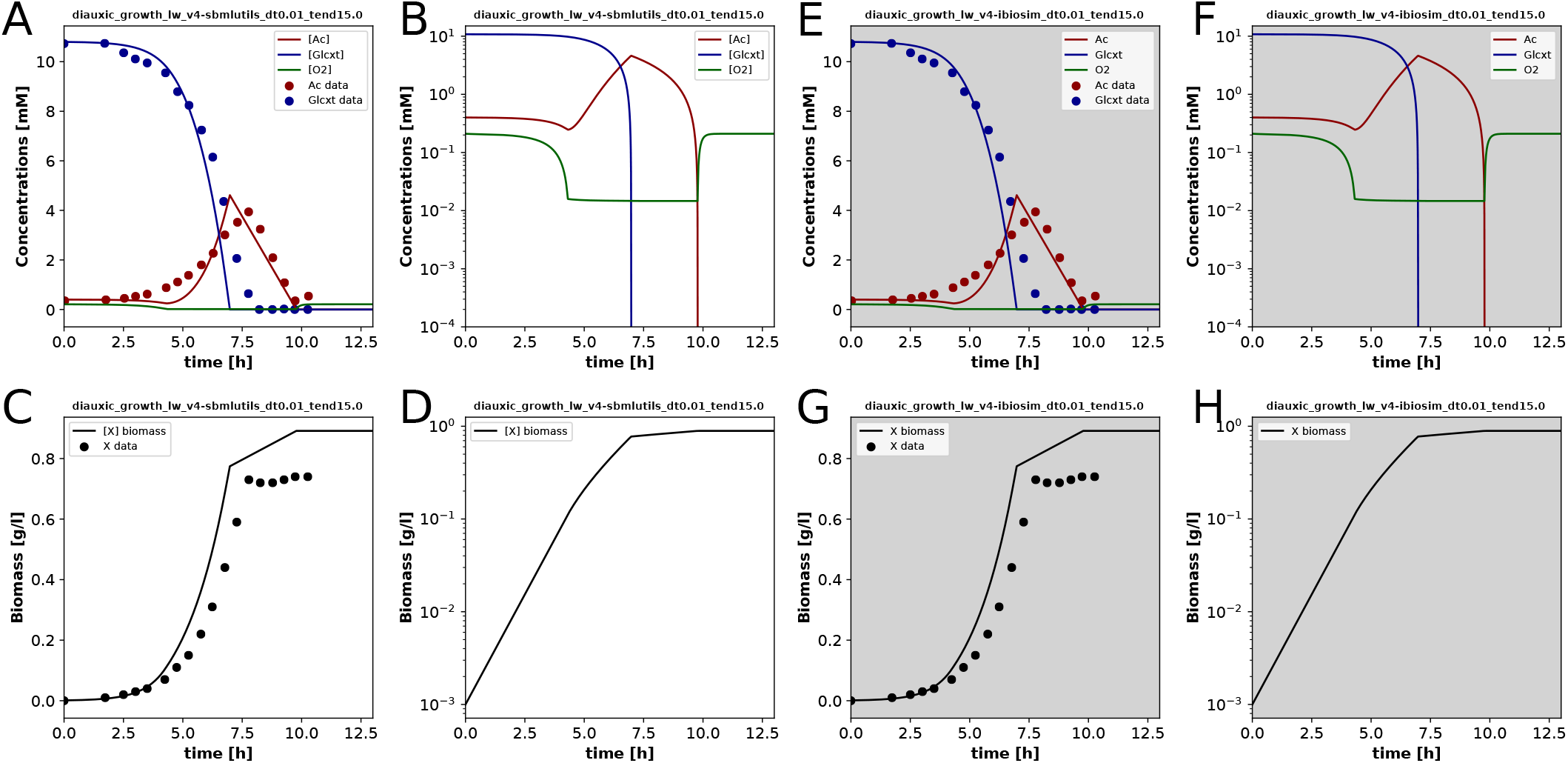
Simulation results for the diauxic growth of *Escherichia coli* in `sbmlutils` (**A, B, C, D**) and `iBioSim` (**E, F, G, H**). The model is able to reproduce the general behavior from experimental data. The cell is growing exponentially while glucose is present, but when the cell runs out of glucose, growth slows down and is limited mainly by oxygen. However, when the cell runs out of glucose and oxygen, growth diminishes significantly. The model was simulated for 15[h] with a time step `dt` of 0.01[h].

We hereby showed that our schema is able to encode published DFBA models, resulting in a reproducible and exchangeable model representation between tools.

### 3.4 *Escherichia coli* Core Metabolism (ecoli)

To demonstrate the feasibility of the proposed schema and method for real-world examples of DFBAs, a larger metabolic network for the core metabolism of *Escherichia coli* (Orth et al., 2010) was encoded in the proposed schema and simulated as shown in Figure 6. The model is available as COMBINE archive in Supplementary Material S3. The FBA submodel was downloaded from BiGG (King et al., 2016) (core metabolism of *Escherichia coli* str. K-12 substr. MG1655) and transformed to an DFBA model in an automatic fashion using `sbmlutils`. BiGG models encode the exchangeable species via annotated exchange reactions which allows an automatic inference of the dynamic species. Only additional information required to run a DFBA simulations are initial concentrations for the species. The automatic encoding of larger scale examples demonstrates the scalability of the proposed encoding approach.

**Figure 6.**
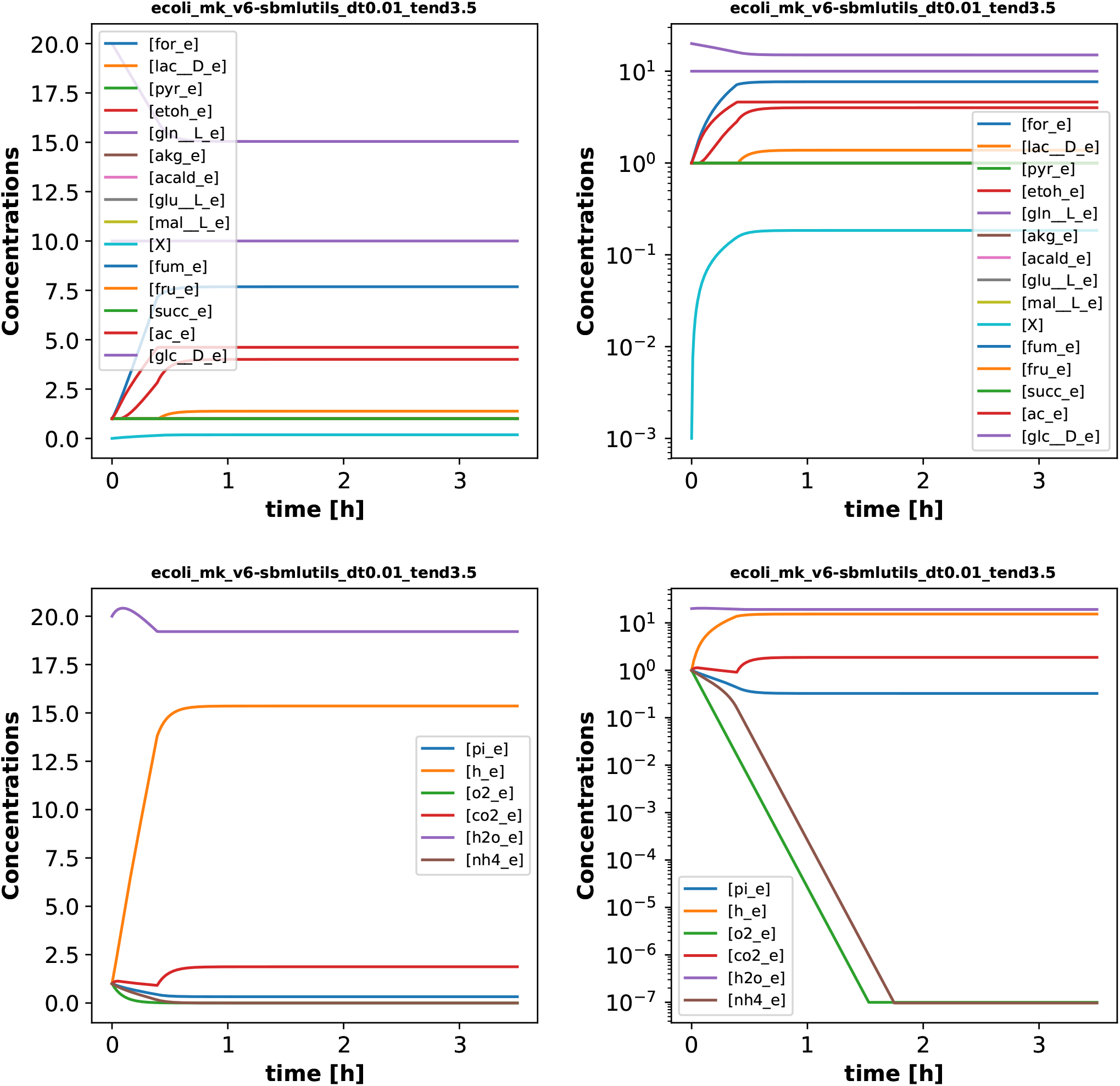
DFBA simulation results for core metabolism of *Escherichia coli* with `sbmlutils`. The proposed approach can be used in larger models, such as the *Escherichia coli* model described in the paper. The model is growing aerobically on glucose in the initial phase and reaches a steady state after oxygen is consumed. The model was simulated for 3.5[h] with a time step `dt` of 0.01[h].

While `sbmlutils` is able to find a solution for the model, `iBioSim` cannot as it runs into an unfeasible solution in the middle of simulation. This captures the well-known problem of DFBA with multiple solutions. The FBA problem is not constrained enough to result in a unique solution and depending on which solution the simulator picks, different solutions and thereby trajectories arise. Despite the existence of multiple solutions, tools and LP solvers typically pick solutions deterministically. Hence, single tools can reproduce their own results, but results are irreproducible between different implementations. Without the use of standards, this could never be demonstrated because variations in results could be due to discrepancies in the model, and not in the tool.

## 4 DISCUSSION

The ability to encode hybrid models, modularity and reproducibility of models are indispensable for encodings of complex models in computational biology. In this work, we presented such an approach, which allows a clear separation of the different modeling formalisms via hierarchical models and defining the interfaces between the submodels. Here we propose and implement an exchangeable and reproducible hybrid modeling scheme. This scheme for encoding DFBA models has been implemented in two different tools, demonstrating the exchangeability and reproducibility of our approach on various examples models. `iBioSim` and `sbmlutils` are freely available for download and offer the necessary infrastructure for anyone to develop DFBA models using the proposed scheme. Currently, the proposed approach supports the modeling of DFBA models based on the SOA simulation algorithm. Hence, our approach only covers a subset of DFBA algorithms and a subset of possible frameworks.

Most DFBA models are stiff. Hence, short time steps are required for stability and for accurate results. Due to the need for short time steps, the SOA approach is computationally expensive. Future directions include the exploration of adaptive time steps for executing the DFBA with SOA, alternative DFBA methods, such as DOA or DA, and extending our scheme to encode such models.

Our current approach is limited to the coupling of ODEs to FBA models. Different hybrid modeling types, such as any mixtures of differential equations, stochastic processes, or Boolean models may yield promising results in the future. The proposed approach of decoupling different modeling formalisms via the comp package could work similarly for other modeling frameworks like Boolean models.

So far, only small to medium-size DFBA models have been encoded in our proposed approach. For future work, we will encode genome-scale metabolic models such as HepatoNet1 (Gille et al., 2010) which will allow us to assess the scalability and performance of the proposed approach.

## Supporting information

Supplemental Material 1 - Schema for encoding DFBA in SBML

Supplemental Material 2 - Reproducibility of results between sbmlutils and iBioSim

Supplemental Material 3 - COMBINE archives and Cytoscape figures for DFBA models.

## AUTHOR CONTRIBUTIONS

MK and LHW designed the study, developed the computational models, implemented and performed the analysis, and wrote the initial draft of the manuscript. All authors discussed the results. All authors contributed to and revised the manuscript critically.

## DATA AVAILABILITY STATEMENT

### Availability

All materials and models are available from https://github.com/matthiaskoenig/dfba. The tools used in this project are freely available: `iBioSim` at https://www.async.ece.utah.edu/ibiosim and `sbmlutils` at https://github.com/matthiaskoenig/sbmlutils/.

## 5 SUPPLEMENTARY MATERIAL

Supplementary Material is available online.

**S1** Schema for encoding DFBA in SBML.

**S2** Reproducibility results between `sbmlutils` and `iBioSim`.

**S3** texttttoy wholecell COMBINE archive and Cytoscape figure for DFBA models (minimal model, minimal glycolysis model, diauxic model, and Escherichia coli core model).

## REFERENCES

Bergmann, F. T., Adams, R., Moodie, S., Cooper, J., Glont, M., Golebiewski, M., et al. (2014). COMBINE archive and OMEX format: One file to share all information to reproduce a modeling project. BMC bioinformatics 15, 369. doi:10.1186/s12859-014-0369-z

Bergmann, F. T., Cooper, J., König, M., Moraru, I., Nickerson, D., Le Novère, N., et al. (2018). Simulation Experiment Description Markup Language (SED-ML) level 1 version 3 (L1V3). Journal of Integrative Bioinformatics 15, /j/jib.2018.15.issue–1/jib–2017–0086/jib–2017–0086.xml. doi:10.1515/jib-2017-0086

Bordbar, A., Monk, J. M., King, Z. A., and Palsson, B. O. (2014). Constraint-based models predict metabolic and associated cellular functions. Nature Reviews. Genetics 15, 107–120. doi:10.1038/nrg3643

Courtot, M., Juty, N., Knüpfer, C., Waltemath, D., Zhukova, A., Dräger, A., et al. (2011). Controlled vocabularies and semantics in systems biology. Molecular Systems Biology 7, 543. doi:10.1038/msb.2011.77

Ebrahim, A., Lerman, J. A., Palsson, B. O., and Hyduke, D. R. (2013). COBRApy: COnstraints-Based Reconstruction and Analysis for Python. BMC systems biology 7, 74. doi:10.1186/1752-0509-7-74

Gille, C., Bölling, C., Hoppe, A., Bulik, S., Hoffmann, S., Hübner, K., et al. (2010). HepatoNet1: A comprehensive metabolic reconstruction of the human hepatocyte for the analysis of liver physiology. Molecular Systems Biology 6, 411. doi:10.1038/msb.2010.62

Gomez, J. A., Höffner, K., and Barton, P. I. (2014). DFBAlab: A fast and reliable MATLAB code for dynamic flux balance analysis. BMC Bioinformatics 15, 409. doi:10.1186/s12859-014-0409-8

Hanly, T. J. and Henson, M. A. (2011). Dynamic flux balance modeling of microbial co-cultures for efficient batch fermentation of glucose and xylose mixtures. Biotechnology and Bioengineering 108, 376–385. doi:10.1002/bit.22954

Hjersted, J. L., Henson, M. A., and Mahadevan, R. (2007). Genome-scale analysis of Saccharomyces cerevisiae metabolism and ethanol production in fed-batch culture. Biotechnology and Bioengineering 97, 1190–1204. doi:10.1002/bit.21332

Höffner, K., Harwood, S. M., and Barton, P. I. (2013). A reliable simulator for dynamic flux balance analysis. Biotechnology and Bioengineering 110, 792–802. doi:10.1002/bit.24748

Hoops, S., Sahle, S., Gauges, R., Lee, C., Pahle, J., Simus, N., et al. (2006). COPASI—a COmplex PAthway SImulator. Bioinformatics 22, 3067–3074. doi:10.1093/bioinformatics/btl485

Hucka, M., Bergmann, F. T., Chaouiya, C., Dräger, A., Hoops, S., Keating, S. M., et al. (2019). The Systems Biology Markup Language (SBML): Language specification for level 3 version 2 core release 2. Journal of Integrative Bioinformatics 16, /j/jib.2019.16.issue–2/jib–2019–0021/jib–2019–0021.xml. doi:10.1515/jib-2019-0021

Hucka, M., Finney, A., Sauro, H. M., Bolouri, H., Doyle, J. C., Kitano, H., et al. (2003). The systems biology markup language (SBML): A medium for representation and exchange of biochemical network models. Bioinformatics (Oxford, England) 19, 524–531. doi:10.1093/bioinformatics/btg015

Karr, J. R., Sanghvi, J. C., Macklin, D. N., Gutschow, M. V., Jacobs, J. M., Bolival, B., et al. (2012). A whole-cell computational model predicts phenotype from genotype. Cell 150, 389–401. doi:10.1016/j.cell.2012.05.044

Kauffman, S. A. (1969). Metabolic stability and epigenesis in randomly constructed genetic nets. Journal of Theoretical Biology 22, 437–467. doi:10.1016/0022-5193(69)90015-0

Keating, S. M., Waltemath, D., König, M., Zhang, F., Dräger, A., Chaouiya, C., et al. (2020). SBML Level 3: An extensible format for the exchange and reuse of biological models. Molecular systems biology 16, e9110. doi:10.15252/msb.20199110

King, Z. A., Lu, J., Dräger, A., Miller, P., Federowicz, S., Lerman, J. A., et al. (2016). BiGG Models: A platform for integrating, standardizing and sharing genome-scale models. Nucleic Acids Research 44, D515–522. doi:10.1093/nar/gkv1049

Kitano, H. (2002). Computational systems biology. Nature 420, 206–210. doi:10.1038/nature01254

[Dataset]König, M. (2022). Sbmlutils: Python utilities for SBML. Zenodo. doi:10.5281/zenodo.6231726

König, M., Dräger, A., and Holzhütter, H.-G. (2012). CySBML: A Cytoscape plugin for SBML. Bioinformatics (Oxford, England) 28, 2402–2403. doi:10.1093/bioinformatics/bts432

Le Novère, N., Hucka, M., Mi, H., Moodie, S., Schreiber, F., Sorokin, A., et al. (2009). The Systems Biology Graphical Notation. Nature Biotechnology 27, 735–741. doi:10.1038/nbt.1558

Lequeux, G., Beauprez, J., Maertens, J., Van Horen, E., Soetaert, W., Vandamme, E., et al. (2010). Dynamic metabolic flux analysis demonstrated on cultures where the limiting substrate is changed from carbon to nitrogen and vice versa. Journal of Biomedicine & Biotechnology 2010, 621645. doi:20160811101003

Luo, R.-Y., Liao, S., Tao, G.-Y., Li, Y.-Y., Zeng, S., Li, Y.-X., et al. (2006). Dynamic analysis of optimality in myocardial energy metabolism under normal and ischemic conditions. Molecular Systems Biology 2, 2006.0031. doi:10.1038/msb4100071

Mahadevan, R., Edwards, J. S., and Doyle, F. J. (2002). Dynamic flux balance analysis of diauxic growth in Escherichia coli. Biophysical Journal 83, 1331–1340. doi:10.1016/S0006-3495(02)73903-9

Mahadevan, R. and Schilling, C. H. (2003). The effects of alternate optimal solutions in constraint-based genome-scale metabolic models. Metabolic Engineering 5, 264–276. doi:10.1016/j.ymben.2003.09.002

Meadows, A. L., Karnik, R., Lam, H., Forestell, S., and Snedecor, B. (2010). Application of dynamic flux balance analysis to an industrial Escherichia coli fermentation. Metabolic Engineering 12, 150–160. doi:10.1016/j.ymben.2009.07.006

Morris, M. K., Saez-Rodriguez, J., Sorger, P. K., and Lauffenburger, D. A. (2010). Logic-based models for the analysis of cell signaling networks. Biochemistry 49, 3216–3224. doi:10.1021/bi902202q

Neal, M. L., Gennari, J. H., Waltemath, D., Nickerson, D. P., and König, M. (2020). Open modeling and exchange (OMEX) metadata specification version 1.0. Journal of Integrative Bioinformatics 17. doi:10.1515/jib-2020-0020

Neal, M. L., König, M., Nickerson, D., Misirli, G., Kalbasi, R., Dräger, A., et al. (2019). Harmonizing semantic annotations for computational models in biology. Briefings in Bioinformatics 20, 540–550. doi:10.1093/bib/bby087

Orth, J. D., Fleming, R. M. T., and Palsson, B. Ø. (2010). Reconstruction and use of microbial metabolic networks: The core Escherichia coli metabolic model as an educational guide. EcoSal Plus 4. doi:10.1128/ecosalplus.10.2.1

Pizarro, F., Varela, C., Martabit, C., Bruno, C., Pérez-Correa, J. R., and Agosin, E. (2007). Coupling kinetic expressions and metabolic networks for predicting wine fermentations. Biotechnology and Bioengineering 98, 986–998. doi:10.1002/bit.21494

Savinell, J. M. and Palsson, B. O. (1992). Network analysis of intermediary metabolism using linear optimization. I. Development of mathematical formalism. Journal of Theoretical Biology 154, 421–454. doi:10.1016/s0022-5193(05)80161-4

Smith, L. P., Hucka, M., Hoops, S., Finney, A., Ginkel, M., Myers, C. J., et al. (2015). SBML Level 3 package: Hierarchical Model Composition, Version 1 Release 3. Journal of Integrative Bioinformatics 12, 268. doi:10.2390/biecoll-jib-2015-268

Somogyi, E. T., Bouteiller, J.-M., Glazier, J. A., König, M., Medley, J. K., Swat, M. H., et al. (2015). libRoadRunner: A high performance SBML simulation and analysis library. Bioinformatics (Oxford, England) 31, 3315–3321. doi:10.1093/bioinformatics/btv363

Thomas, R. (1973). Boolean formalization of genetic control circuits. Journal of Theoretical Biology 42, 563–585. doi:10.1016/0022-5193(73)90247-6

Tomita, M., Hashimoto, K., Takahashi, K., Shimizu, T. S., Matsuzaki, Y., Miyoshi, F., et al. (1999). E-CELL: Software environment for whole-cell simulation. Bioinformatics (Oxford, England) 15, 72–84. doi:10.1093/bioinformatics/15.1.72

Varma, A., Boesch, B. W., and Palsson, B. O. (1993). Biochemical production capabilities of Escherichia coli. Biotechnology and Bioengineering 42, 59–73. doi:10.1002/bit.260420109

Varma, A. and Palsson, B. O. (1994). Stoichiometric flux balance models quantitatively predict growth and metabolic by-product secretion in wild-type Escherichia coli W3110. Applied and Environmental Microbiology 60, 3724–3731. doi:10.1128/aem.60.10.3724-3731.1994

Waltemath, D., Karr, J. R., Bergmann, F. T., Chelliah, V., Hucka, M., Krantz, M., et al. (2016). Toward community standards and software for whole-cell modeling. IEEE transactions on bio-medical engineering 63, 2007–2014. doi:10.1109/TBME.2016.2560762

Watanabe, L., Nguyen, T., Zhang, M., Zundel, Z., Zhang, Z., Madsen, C., et al. (2019). iBioSim 3: A tool for model-based genetic circuit design. ACS synthetic biology 8, 1560–1563. doi:10.1021/acssynbio.8b00078

Watanabe, L. H. and Myers, C. J. (2014). Hierarchical stochastic simulation algorithm for sbml models of genetic circuits. Frontiers in Bioengineering and Biotechnology 2, 55. doi:10.3389/fbioe.2014.00055

Zhukova, A., Waltemath, D., Juty, N., Laibe, C., and Le Novère, N. (2011). Kinetic simulation algorithm ontology. Nature Precedings, 1–1doi:10.1038/npre.2011.6330.1

