## Supplemental Material 1 - Schema for encoding DFBA in SBML for "Dynamic Flux Balance Analysis Models in SBML"

### Guidelines for encoding DFBA models in SBML

**version: 0.3.2**

### Introduction

This document describes rules and guidelines for encoding Dynamic Flux Balance Analysis (DFBA) models in the Systems Biology Markup Language (SBML), a free and open interchange format for computer models of biological processes.

Note that these guidelines have been proposed by iBioSim or sbmlutils as ground rules for encoding and simulating DFBA models in these tools in an interchangeable and reproducible manner. It is by no means a community agreement. However, we highly encourage everyone who wants to encode DFBA models and tool developers to follow these guidelines.

The document is structured into the following sections

- **Section A**: describes how to encode DFBA models in SBML.
- **Section B**: provides information on how simulators should execute models provided in the format of Section A).
- **Section C**: provides answers to frequently asked questions.

DFBA Implementation based on these rules and guidelines are provided by iBioSim or sbmlutils. All supplementary information, including the latest version of this document as well as example models implementing the DFBA rules and guidelines, are provided in at https://github.com/matthiaskoenig/dfba.

The rules and guidelines for DFBA encoding were developed for models using the stationary optimization approach (SOA).

We expect readers to be familiar with the concepts of SBML and the `fbc` and `comp` packages and refer to the respective specifications http://sbml.org/Documents/Specifications for additional information. Also we expect readers to be familiar with the concepts of DFBA and refer to the respective literature.

The following conventions are used throughout this document:

- Required rules are stated via **MUST**, i.e., DFBA models in SBML must implement these rules.
- Guidelines which are recommended to be followed are indicated by **SHOULD**, i.e., it is good practice to follow these guidelines, but they are not required for an executable and reproducible DFBA model encoded in SBML. The provided implmentations by iBioSim and sbmlutils will run the DFBA even if these recommendations are not followed.
- Additional information for clarification is provided by **CAN**, i.e., it is clarified that this is allowed to remove ambiguities.

- Curly brackets, i.e, `{ }` functions as place holders which are instantiated with actual information. For instance the reaction id `{rid}` means that `{rid}` is replaced with the actual id of the reaction.

##### List of abbreviations

The following abbreviations are used in this document

- DFBA - Dynamic Flux Balance Analysis
- FBA -Flux Balance Analysis
- SBML - Systems Biology Markup Language
- SOA - Stationary optimization approach

### A Encoding DFBA models in SBML

This section describes rules and guidelines for encoding DFBA models in SBML. The proposed schema uses SBML `core`, SBML `comp` for model compositions, and SBML `fbc` to encode FBA related information.

The core concept behind this guidelines and rules is to encode models with different modeling frameworks, i.e., kinetic models and FBA models, as well as models with different functions, i.e., updating or calculation of flux bounds within separate submodels. These submodels are connected into the overall DFBA model using hierarchical model composition based on `comp`.

Two main links are required between the FBA model and the kinetic models:

- Update of flux bounds in the FBA model from the kinetic model
- Update of reaction fluxes in the kinetic model from the FBA solution

The DFBA models consists of different components performing parts of the DFBA task:

- `TOP` model: DFBA comp model that includes all submodels and their corresponding connections. The `TOP` model is the main SBML model, containing the other submodels. The `TOP` model encodes the kinetic model parts of the DFBA (besides bounds calculation and updates from FBA).
- `KINETIC` submodel: kinetic part of the DFBA model
- `FBA` submodel: FBA part of the DFBA model. The `FBA` model defines the FBA submodel using the `fbc` package.
- `BOUNDS` submodel: calculation of the upper and lower bounds for the `FBA` model. The `BOUNDS` model defines all logic for the update of the FBA bounds.
- `UPDATE` submodel: calculation of the updated `KINETIC` part from the `FBA` solution

An overview of the different submodels and their connections is provided in the following diagram:

#### A.1 `DFBA` model

In this subsection general rules and guidelines are defined.

- `[DFBA-R0001]` The DFBA model **MUST** be a single SBML `comp` model.
- `[DFBA-R0002]` The DFBA submodels **MUST** be encoded in the DFBA model via `comp:SubModels`.
- `[DFBA-R0003]` The DFBA submodels **MUST** be defined via `comp:ExternalModelDefinitions`.
- `[DFBA-R0004]` The DFBA model and all submodels **MUST** be encoded in `SBML L3V1` or higher.
- `[DFBA-R0005]` The DFBA model and all submodels **MUST** be valid SBML.
- `[DFBA-R0006]` The DFBA model **MUST** be encoded using SBML `core` and the SBML packages `comp` and `fbc`.
- `[DFBA-R0007]` The DFBA model **MUST** consist of the `TOP` model and at least three submodels, the required`FBA`, `BOUNDS` and `UPDATE` submodel.
- `[DFBA-G0001]` The model and submodels **SHOULD** contain their respective function in the `model id`, `model name` and `filename`, i.e. the strings `TOP` or `top`, `FBA` or `fba`, `BOUNDS` or `bounds`, and `UPDATE` or `update`.
- `[DFBA-G0002]` The SBOTerms on the `submodel` object **SHOULD** be identical to the SBOTerm on the `Model` object of all submodels.
- The `TOP` model **CAN** contain additional submodels.
- The DFBA model and all submodels **CAN** have additional packages than `fbc` and `comp`.

##### fbc

- `[DFBA-R0008]` There **MUST** exist exactly one submodel with the `fbc` package and the SBOTerm `SBO:0000624` (flux balance framework) on the `model` element. This model is called the `FBA` submodel for the `DFBA`.
- `[DFBA-R0009]` The `FBA` submodel **MUST** be encoded using `fbc-v2` with `strict=true`.
- There **CAN** be other submodels with the `fbc` package, but not with the SBOTerm `SBO:0000624` (flux balance framework) on the model element. These submodels **CAN** be either `strict=True` or `strict=False`.

##### ports

Objects in the different submodels are linked via `comp:Ports`.

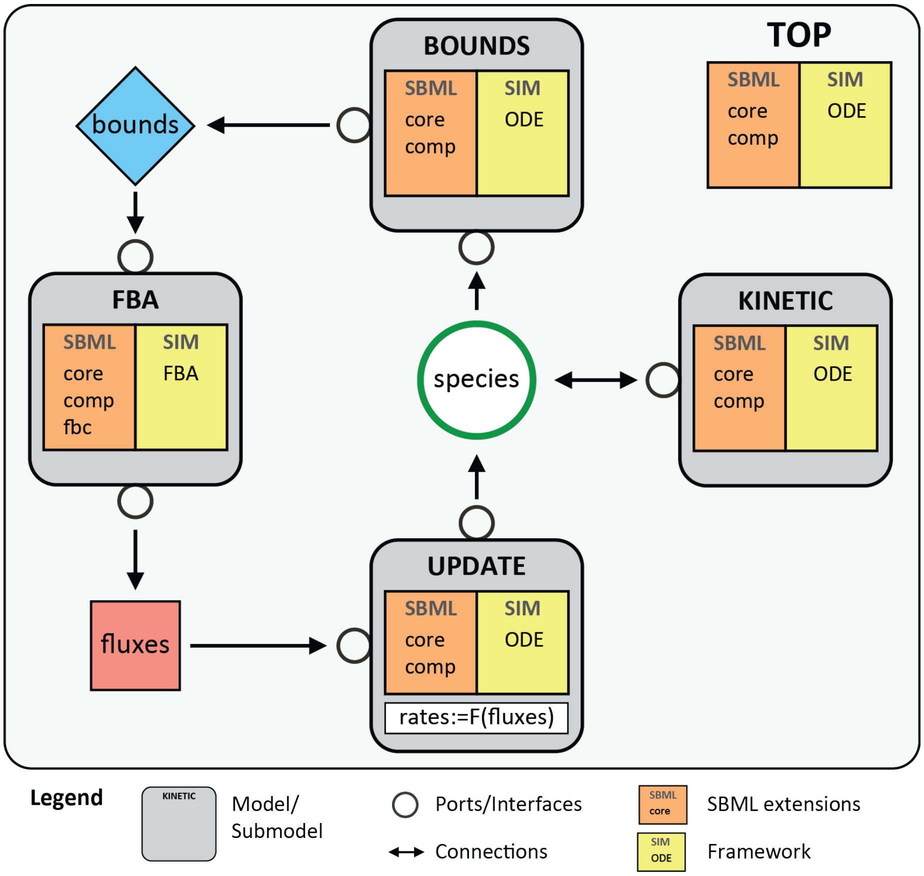


Figure 1: Overview of schema for encoding DFBA models in SBML. The hierarchical SBML model is composed of a top-level model with four submodels: FBA, BOUNDS, UPDATE, and KINETIC. The individual submodels are connected via ports. The respective SBML packages used are listed in the models, as well as the simulation framework used. The BOUNDS submodel calculates the upper and lower flux bounds based on metabolite availability. The FBA submodel computes the reaction fluxes of the metabolic fbc model using the bounds as constraints. The UPDATE submodel calculates the dynamic update of the dynamic metabolites affected by the FBA model. The rates of change are herby functions of the FBA fluxes. The KINETIC submodel includes all of the other processes in the model, which may affect or be affected by entities in metabolism. The top-level model ties together the different submodels using SBML comp replacements and replacedBy constructs.

- [DFBA-R0010] All ReplacedBy and Replacements **MUST** be done via ports which are identified via idRef.
- [DFBA-R0011] Objects which are linked via ports in the different submodels **MUST** have identical ids in the the different submodels.
- [DFBA-R0012] In addition, the respective ports of the linked objects **MUST** have the same ids.
- [DFBA-G0003] All Ports **SHOULD** have the id {idRef}_port for an object with idRef={idRef}.

###### units

- [DFBA-G0004] All models **SHOULD** contain units.
- [DFBA-G0005] The units of the submodel **SHOULD** be identical and be replaced by the top model.

##### A.2 TOP model

In this subsection the rules and guidelines for the TOP model are defined.

- [TOP-R0001] The TOP model **MUST** have the SBOTerm SBO:0000293 (non-spatial continuous framework) on the Model element.
- [TOP-R0002] The TOP model **MUST** have exactly one submodel with the SBOTerm SBO:0000624 (flux balance framework) on the Model element.

#### dt

- [TOP-R0003] The TOP DFBA model **MUST** contain a parameter dt which defines the step size of the FBA optimizations, i.e. after which time interval the FBA is performed.
- [TOP-R0004] The dt parameter **MUST** be annotated with the SBOTerm SBO:0000346 (temporal measure).
- [TOP-R0021] The dt Parameter **MUST** be constant.
- [TOP-R0005] If the dt parameter has units, than they **MUST** be identical to the timeUnits of the model.

###### dummy species & exchange reactions

Dummy species are required for the definition of dummy reactions in SBML L3V1, because every reaction requires at least one reactant or product (the following rules can be relaxed in SBML L3V2).

- [TOP-R0006] The top model **MUST** have a dummy species with id="dummy_S".

- `[TOP-R0007]` For every exchange reaction in the `FBA` submodel, there **MUST** exist a dummy exchange reaction in the `TOP` model.
- `[TOP-R0008]` Each dummy exchange reaction **MUST** include the dummy species `dummy_S` as product with stochiometry `1.0`.
- `[TOP-R0009]` The dummy exchange reaction **MUST NOT** have any other reactants, products or modifiers than `dummy_S`, i.e. `-> dummy_S`
- `[TOP-G0001]` The id of the dummy reaction **SHOULD** be identical to the respective exchange reaction, i.e. `id="{rid}"` for the exchange reaction with `id="{rid}"` in the `FBA` submodel.
- `[TOP-G0002]` The dummy species **SHOULD NOT** have and `compartment` set.
- `[TOP-G0003]` The dummy species **SHOULD** have the SBOTerm `SBO:0000291` (empty set).
- `[TOP-G0004]` The dummy reactions **SHOULD** have the SBOTerm `SBO:0000631` (pseudoreaction).
- `[TOP-G0005]` The dummy species **CAN** be in an arbitrary `compartment` of the `TOP` model.

##### exchange species

- `[TOP-R0010]` The `TOP` model **MUST** contain a species for every species which has an exchange reaction in the `FBA` model (exchange `species`).
- `[TOP-R0011]` The exchange `species` **MUST** replace the corresponding species in the `UPDATE` and `BOUNDS` model via `ReplacedElements`.

##### flux parameters & flux assignmentRules

- `[TOP-R0012]` For every dummy `Reaction` in the `TOP` model, a corresponding flux `Parameter` **MUST** exist in the `TOP` model which is `constant=true` with the id `{pid}`.
- `[TOP-R0013]` For every dummy exchange `Reaction` with `id={rid}` and corresponding flux `Parameter` with `id={pid}` in the top model an `AssignmentRule` in the `TOP` model **MUST** exist of the form `{pid} = {rid}`.
- `[TOP-G0005]` The flux parameter **SHOULD** have the id `p{rid}` for the corresponding dummy reaction `{dummy_rid}`, e.g. `pEX_Glc` for `EX_Glc`.
- `[TOP-G0006]` The flux `Parameters` **SHOULD** have the SBOTerm `SBO:0000612` (rate of reaction).
- `[TOP-G0007]` The flux `AssignmentRules` **SHOULD** have the SBOTerm `SBO:0000391` (steady state expression).

##### replacedBy

- [TOP-R0014] Every dummy reaction in the TOP model with id="dummy_{rid}" **MUST** be replaced via a comp:ReplacedBy with the corresponding exchange reaction with id={EX_rid} from the FBA submodel. The comp:ReplacedBy uses the portRef of the exchange reaction {EX_rid}_port.

These replacements update the ODE fluxes in the TOP model by replacing the dummy Reaction by the corresponding FBA reaction.

##### replacements

- [TOP-R0015] For every parameter that is used as a flux bound, other than default ones, for a reaction in the FBA submodel, there **MUST** be a replacing parameter in the TOP model.
- [TOP-R0016] For the dt parameter in the BOUNDS model there must be a replacement with the TOP dt parameter.
- [TOP-R0017] For every species that is used for bounds calculation in the BOUNDS model (this includes all exchange species) there **MUST** exist a replacement species in the TOP model.
- [TOP-R0018] For every species that is updated in the UPDATE models there **MUST** exist a replacement species in the TOP model.
- [TOP-R0019] TODO:For every uper and lower bound parameter ... (exchange reactions & kinetic reactions)
- [TOP-G0008] The replaced species in the BOUNDS and UPDATE submodels **SHOULD** be connected via the same replacing species in the TOP model.

#### A.3 FBA submodel

In this subsection the rules and guidelines for the FBA model are defined. The FBA model defines the FBA submodel using the fbc package.

- [FBA-R0001] The Model element of the FBA submodel **MUST** have the SBOTerm SBO:0000624 (flux balance framework) on the Model element.
- [FBA-R0002] The FBA model **MUST** be encoded using the SBML package fbc-v2 with strict=true.
- [FBA-R0003] The reactions in the FBA model **MUST NOT** have any KineticLaw.

##### objective function

- [FBA-R0004] The FBA model **MUST** contain at least one objective function. Objective functions **CAN** be maximize or minimize.

- `[FBA-R0005]` The objective function for the DFBA model **MUST** be the active objective in the `FBA` model.

##### exchange reactions

Unbalanced species in the `FBA` model correspond to species in the kinetic model which are changed via the FBA fluxes. Unbalanced species are encoded by the means of exchange reactions.

- `[FBA-R0006]` Unbalanced `species` in the FBA **MUST** be encoded by creating an exchange reaction for the respective species.
- `[FBA-R0007]` The exchange `Reactions` **MUST** have the `Species` which is changed by the reaction (unbalanced `Species` in FBA) as substrate with stoichiometry `1.0` and have no products, i.e. have the form `1.0 {sid} ->` with `{sid}` being the `Species` id.
- `[FBA-G0001]` The exchange `Reactions` **SHOULD** have the SBOterm `SBO:0000627` (exchange reaction).
- `[FBA-G0002]` The exchange `Reactions` **SHOULD** be named `EX_{sid}`, i.e. consist of the prefix `EX_` and the `Species` id `{sid}`.
- `[FBA-G0003]` Exchange reactions **SHOULD NOT** have a `compartment`.

##### boundaryCondition

- `[FBA-R0009]` All `Species` in the FBA model **MUST** have `boundaryCondition=False`.

##### reaction flux bounds

- `[FBA-R0010]` All exchange reactions **MUST** have individual `Parameters` for the upper and lower bound which are not used by other reactions (unless using default bounds).
- `[FBA-G0004]` The `Parameters` for the upper and lower bounds of reactions **SHOULD** have the ids `ub_{rid}` and `lb_{rid}` with `{rid}` being the respective reaction id.
- `[FBA-G0005]` The `Parameters` describing the flux bounds **SHOULD** have the SBOTerm `SBO:0000625` (flux bound).

##### ports

- `[FBA-R0011]` All exchange reactions **MUST** have a port.
- `[FBA-R0012]` All upper and lower bounds of exchange reactions **MUST** have a port.

#### A.4 BOUNDS submodel

In this subsection the rules and guidelines for the `BOUNDS` model are defined. The `BOUNDS` submodel calculates the upper and lower bounds for the `FBA` model. For this calculation the `Species` changed via exchange `Reactions` in the FBA and the time step `dt` are required.

The parameter `dt` is used in calculating the upper and lower bounds based on the availability of the species in the exchange `Reactions`. This ensures that the FBA solution cannot take more than the available species amounts in the time step of duration `dt` and is consistent for the time step with the available resources.

- `[BND-R0001]` The `BOUNDS` model **MUST** have the SBOTerm `SBO:0000293` (non-spatial continuous framework) on the `Model` element.

### dt

- `[BND-R0002]` The `BOUNDS` model **MUST** contain the parameter `dt` which defines the step size of the FBA optimizations.
- `[BND-R0016]` The `dt` Parameter **MUST** be constant.
- `[BND-R0004]` The `dt` parameter **MUST** be annotated with the SBOTerm `SBO:0000346` (temporal measure).

##### bounds species & assignment rules

- `[BND-R0005]` The `BOUNDS` submodel **MUST** contain all exchange `Species`, i.e. `Species` which are reactants in `FBA` exchange `Reactions`.
- `[BND-R0006]` The `BOUNDS` submodel **MUST** contain all `Compartments` which are used in exchange `Species`.
- `[BND-R0007]` The `BOUNDS` model **MUST** contain `Parameters` for all upper and lower flux bounds of exchange `Reactions`.
- `[BND-R0008]` The `BOUNDS` model **MUST** contain `FunctionDefinitions` for `min` and `max` of the form
  `min=lambda( x,y, piecewise(x,lt(x,y),y) )`
  and
  `max=lambda( x,y, piecewise(x,gt(x,y),y) )`.
- `[BND-R0009]` The `BOUNDS` model **MUST** contain `AssignmentRules` for the update of lower bounds of the exchange reactions of the form `lb_EX_{sid}=max(lb_default, -{sid}*{cid}/dt)` with `{cid}` being the compartment of the species `{sid}`. This ensures that in the time step `dt` not more than the available amounts of the species are used in the `FBA` solution.

- [BND-R0010] If there are additional kinetic lower bounds on the exchange reactions these kinetic bounds **MUST** be used for restricting the bounds via `lb_EX_{sid}=max(lb_kinetic, -{sid}*{cid}/dt)`
- [BND-R0011] The BOUNDS model **MUST** contain the necessary parameter and assignment rules for the update of additional upper and lower bounds of reactions in the FBA which are not exchange reactions. E.g. if there is a time dependent change in an upper bound of an FBA reaction this belongs in the BOUNDS model.
- [BND-G0001] The Parameters describing the flux bounds **SHOULD** have the SBOTerm SBO:0000625 (flux bound).
- The BOUNDS submodel **CAN** calculate additional kinetic bounds for exchange reactions via AssignmentRules, RateRules or EventAssignments.

###### ports

- [BND-R0003] The dt Parameter **MUST** have a Port.
- [BND-R0012] All bound Species used in the BOUNDS model **MUST** have a Port.
- [BND-R0013] All Compartments of bound Species **MUST** have a Port.
- [BND-R0014] All upper and lower bounds of exchange reactions **MUST** have a Port.
- [BND-R0015] All additional kinetic bounds parameter changed in the BOUNDS model **MUST** have a Port.

###### replacedElements

- [BND-R0016] The TOP model **MUST** contain parameters with ReplacedElements for all upper and lower bounds which are changed via the BOUNDS submodel.

##### A.5 UPDATE submodel

In this subsection the rules and guidelines for the UPDATE model are defined. The update submodel performs the update of the species which are changed by the FBA, i.e. the species which have exchange reactions.

- [UPD-R0001] The UPDATE model **MUST** have the SBOTerm SBO:0000293 (non-spatial continuous framework) on the Model element.
- [UPD-R0002] The UPDATE model **MUST** contain corresponding dynamic Species for all Species which are reactants in FBA exchange Reactions.
- [UPD-R0003] The UPDATE model **MUST** contain corresponding compartments for all Species which are reactants in FBA exchange Reactions.

- `[UPD-G0001]` The species in the `UPDATE` submodel **SHOULD** have identical ids to the species in the `FBA` submodel.

###### update reactions & flux parameters

- `[UPD-R0004]` For every `FBA` exchange reaction with id `{rid}` the `UPDATE` model **MUST** contain a respective flux parameter with id `{pid}`.
- `[UPD-R0005]` The every flux parameter in the `UPDATE` submodel the `TOP` model **MUST** have a corresponding flux parameter with a `replacedElement` for the flux parameter in the `UPDATE` model.
- `[UPD-R0006]` For every `FBA` exchange `Reaction` the `UPDATE` model **MUST** contain an update `reaction` with identical reaction equation than the corresponding exchange reaction, i.e. `S ->`.
- `[UPD-R0007]` The update reaction **MUST** have a `KineticLaw` which depends on the flux parameter `{pid_S}` `f(pid_S)` for the `Species` S being updated. In the simplest case the update is performed via `update_S = -pid_S` i.e., the resulting change in Species via the update reaction is than `dS/dt = -pid_S`.
- `[UPD-G0002]` The update reactions **SHOULD** have the SBOTerm `SBO:0000631` (pseudoreaction).
- `[UPD-G0003]` The flux parameters **SHOULD** have the SBOTerm `SBO:0000613` (reaction parameter).
- `[UPD-G0004]` The update reactions **SHOULD** have no `Compartment` set.
- `[UPD-G0005]` The update `Reactions` **SHOULD** have ids of the form `update_{sid}` with `{sid}` being the id of the `Species` which is updated.
- `[UPD-G0006]` The flux `Parameters` in the `UPDATE` model **SHOULD** have identical ids to the flux parameters in the top model.

###### ports

- `[UPD-R0008]` All `Species` used in the `UPDATE` model **MUST** have a port.
- `[UPD-R0009]` All `Compartments` of bound `Species` **MUST** have a port.
- `[UPD-R0010]` All flux `Parameters` **MUST** have a port.

#### A.6 SED-ML and COMBINE archive

DFBA models **SHOULD** be exchanged as COMBINE archives containing all SBML submodels. A simulation experiment description for the DFBA simulation **SHOULD** be provdided in (SED-ML) in the COMBINE archive demonstrating core behavior of the DFBA model, i.e., simple timecourse simulations. In the SED-ML the simulation algorithm **MUST** be provided with the simulation algorithm being from the subset of KISAO terms

- KISAO:0000499 dynamic flux balance analysis (DFBA)

– KISAO:0000500 static optimization approach dynamic flux balance analysis (SOA-DFBA)

The examples and implementations are all based on the static optimization approach (SOA-DFBA).

### B Model Simulation

In this section we describe how models in the DFBA SBML formalism described in section A should be simulated by software. The described simulation and update strategy was implemented in two DFBA simulators: iBioSim and sbmlutils.

#### Static Optimization Approach (SOA)

The DFBA models are solved via a **Static Optimization Approach (SOA)**. The total simulation time is divided into time intervals of length `dt` with the instantaneous optimization (FBA) solved at the beginning of every time interval. The dynamic equations are than integrated over the time interval assuming that the fluxes are constant over the interval. Before every optimization of the FBA part optimization constraints have to be updated from the dynamic part, after every optimization the dynamic variables corresponding to the FBA fluxes have to be updated.

#### Simulation Algorithm

The simulation algorithm starts off by computing the reaction fluxes in the FBA submodel. The reaction fluxes updates the reaction values in the TOP model, which are used to compute the reaction rates in the UPDATE submodel. Once the reaction fluxes are computed by FBA, all NON-FBA submodels are updated concurrently.

```
time = 0
# calculate initial flux bounds
calculate_initial_state()
while (time <= tend){
    # FBA
    set_bounds_fba()
    v_optimal = optimize_fba()

    # ODE
    update_fluxes_ode(v_optimal)
    integrate_ode(start=time, end=time+dt, steps=1)
```

```
    # next time step
    time = time + dt
}
```

- The output time points **MUST** be in agreement with the `dt` parameter, i.e. the interval between subsequent time points **MUST** be `dt`. This does not affect the internal steps of the kinetic solver.
- The model simulation **MUST** abort if the FBA LP probelm is infeasible.
- If the kinetic simulation encounters problems like unfulfilled tolerances the simulation **MUST** stop.
- The flux bounds **MUST** be updated from the kinetic model before the FBA optimization is run.
- The fluxes in the kinetic model **MUST** be set before the kinetic simulation is run.
- For the execution of the kinetic models the comp model is flattend and the flattened model is simulated.

###### FBA optimization

- For the FBA optimization the `reversible` attribute of `Reactions` does not influence the fba solution, Only the upper and lower bounds restrict the possible direction of flux for a reaction.
- The FBA optimization is performed using pFBA (parsimonous FBA) resulting in a Flux distribution with minimal total flux.

##### Tolerances

For the DFBA simulation absolute tolerances `absTol` and `relTol` are defined. These tolerances are used for the kinetic integration. In addition `absTol` is used in the update of the bounds. If the updated bounds are smaller than the absolute tolerance the bounds are set to zero (this avoids infeasible LP problems due to very small negative upper bounds or positive lower bounds).

```
if abs(bound_updated)<= absTol:
    bound_updated = 0
```

#### C Frequently asked questions (FAQ)

##### Are multiple kinetic models supported?

Yes, multiple kinetic submodels can exist in the DFBA. During the kinetic integrations the flattend kinetic model is integrated. However, kinetic submodels **SHOULD** be kept inside the KINETIC submodel.

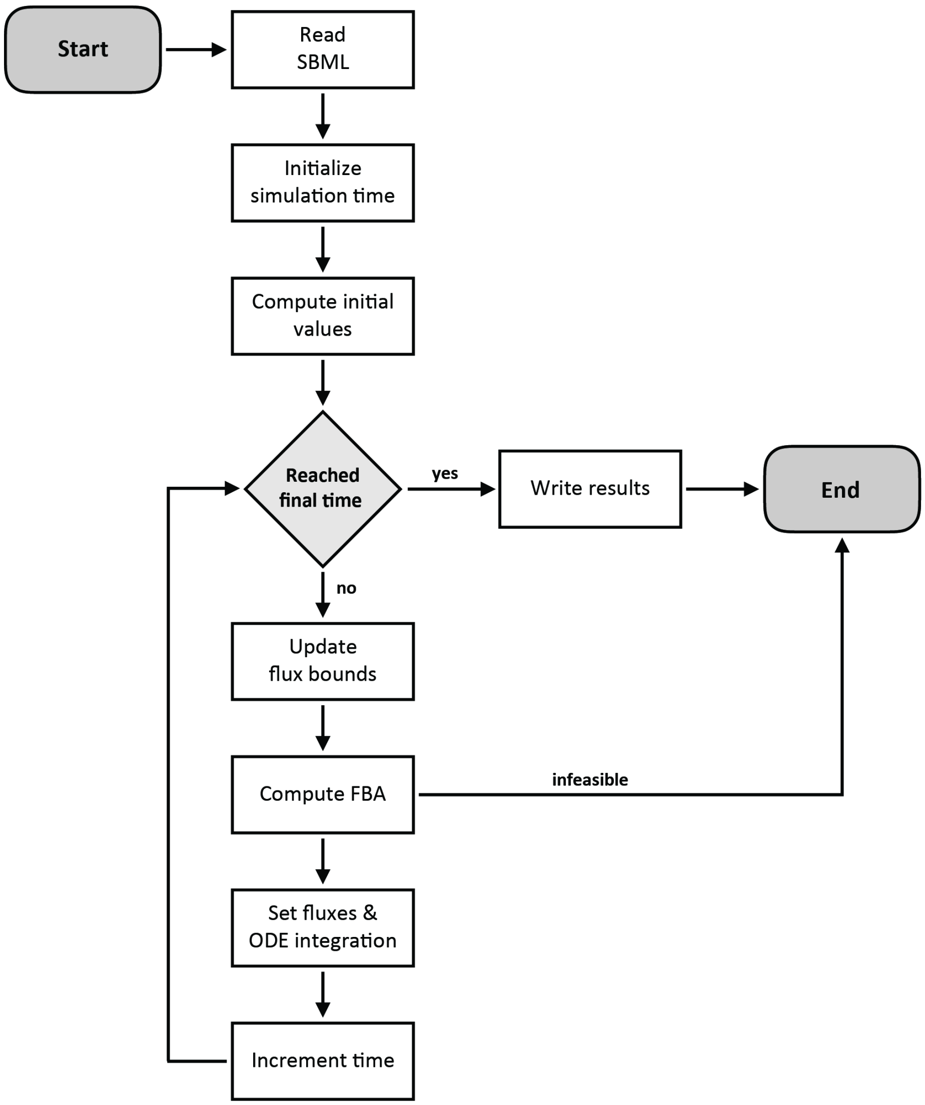


Figure 2: Overview SOA-DFBA simulation algorithm

#### Are multiple FBA submodels supported?

No, in the first version only a single FBA submodel is allowed.

#### Are stochastic & logical models supported?

No, in the first version of the DFBA guidelines and implementation only deterministic kinetic models can be coupled to FBA models. In future versions the coupling of stochastic and/or logical models can be supported. It is possible to encode SBML models with additional modeling frameworks than FBA or deterministic ODE models. Examples are logical models encoded with the SBML package `qual` or stochastic models, i.e. stochastic ODE models. Such models will be considered in future versions.

#### Are variable step sizes supported?

No, currently only fixed step sizes are supported. The simulation steps must be in agreement with the `dt` parameter for bound updates.

#### What SBML constructs are supported by the simulators?

Currently, in `iBioSim` and `sbmlutils` all SBML core constructs are supported in the kinetic models with the exception of `Delay` and `AlgebraicRule`.

#### I am a tool developer and have different ideas about DFBA encoding in SBML. How can I contribute?

You can make suggestions on the Github Issue Tracker. Note this does not guarantee that your suggestions will be adopted. However, we welcome good ideas that would improve our proposed data model idea.

#### What if the `FBA` model has species with `boundaryCondition=True`?

FBA models containing species with `boundaryCondition=True` can easily be converted in supported `FBA` models by setting `boundaryCondition=False` and adding a exchange `Reaction` for the corresponding `Species`.
