## Supplemental Material 2 - Reproducibility of results between sbmlutils and iBioSim for "Dynamic Flux Balance Analysis Models in SBML"

### compare

November 12, 2017

#### 1 Reproducibility between tools

The following scripts checks that solutions are reproducibile between tools. Reproducibility of the model simulations was tested by comparing the numerical SOA results between `sbmlutils` and `iBioSim` for models with unique solutions. Results were assumed as numerical identical if the absolute difference for every time point $t_k$ for all dynamical FBA species $c_k$ in the model was smaller than the tolerance $\epsilon = 1E{-}5$, i.e.,

$$abs(c_i(t_k)_{sbmlutils} - c_i(t_k)_{ibiosim}) \leq \epsilon \; \forall c_i, t_k$$

In [1]:
```
"""
Helper class for comparing simulation results.
"""
import pandas as pd
from matplotlib import pyplot as plt
from pprint import pprint
import warnings

class DataSetsComparison(object):
    """ Comparing two simulation results.
    Currently only supports comparison between two datasets.
    """
    eps = 1E-5  # tolerance for comparison

    def __init__(self, files, dfs, columns=None):
        self.files = files
        self.columns = columns
        for df in dfs:
            # check that identical number of timepoints
            if len(df) != len(dfs[0]):
                raise ValueError("DataFrames have different length: \
                                 {} != {}".format(len(df), len(dfs[0])))

        if columns:
            assert len(self.files) == len(self.columns)
            for column in self.columns:
                assert len(column) == len(self.columns[0])
```

```python
        self.read_dfs(dfs)
        self.diff = self.df_diff()

    def read_dfs(self, dfs):
        """ Read the dataframes using the files and given column ids."""
        self.dfs = []

        for k, df in enumerate(dfs):
            file = self.files[k]
            if self.columns:
                cols = self.columns[k]
                try:
                    df1 = df[cols]
                except KeyError:
                    pprint(df.columns)
                    raise
                df1.columns = self.columns[0]  # unify columns
                self.dfs.append(df1)
            else:
                # no columns specified, necessary to figure out the mapping
                print("-"*40)
                print(file)
                print("-"*40)
                pprint(df.columns)

        return self.dfs

    def df_diff(self):
        """ DataFrame of all differences between the files."""
        return self.dfs[0]-self.dfs[1]

    def is_equal(self):
        """ Check if DataFrames are identical within numerical tolerance."""
        return abs(self.diff.abs().max().max()) <= DataSetsComparison.eps


    def info(self):
        pprint(self.files)
        pprint(self.columns)

    def print_diff(self):
        print("\n# Elements")
        print(self.diff.shape)

        print("\n# Maximum column difference")
        print(self.diff.abs().max())
```

```
            print("\n# Maximum element difference")
            print(self.diff.abs().max().max())

            print("\n# Datasets are equal (diff <= eps={})".format(self.eps))
            print(self.is_equal())
            if not (self.is_equal()):
                warnings.warn("Datasets are not equal !")


        def plot_diff(self):
            for cid in self.diff.columns:
                plt.plot(self.diff[cid], label=cid)
            plt.legend()
            plt.show()

        def report(self):
            print("*" * 80)
            self.info()
            self.print_diff()
            print("*" * 80)
            self.plot_diff()
            self.diff
```

#### 1.1 toy_wholecell

```
In [2]: wholecell_version = 14

        filesets = [
            [
                "./toy_wholecell/toy_wholecell_mk_v14-sbmlutils_dt0.1_tend50.0.csv",
                "./toy_wholecell/toy_wholecell_mk_v14-ibiosim_dt0.1_tend50.0.csv"
            ],
            [
                "./toy_wholecell/toy_wholecell_mk_v14-sbmlutils_dt1.0_tend50.0.csv",
                "./toy_wholecell/toy_wholecell_mk_v14-ibiosim_dt1.0_tend50.0.csv"
            ],
            [
                "./toy_wholecell/toy_wholecell_mk_v14-sbmlutils_dt5.0_tend50.0.csv",
                "./toy_wholecell/toy_wholecell_mk_v14-ibiosim_dt5.0_tend50.0.csv"
            ],
        ]
        for files in filesets:
            wholecell_dsc = DataSetsComparison(
                files=files,
                dfs=[pd.read_csv(file) for file in files],
                columns = [
                    ["time", "[A]", "[C]", "[D]"],
                    ["time", "A", "C", "D"],
```

```
        ]
    )

    wholecell_dsc.report()
```

```
********************************************************************************
['./toy_wholecell/toy_wholecell_mk_v14-sbmlutils_dt0.1_tend50.0.csv',
 './toy_wholecell/toy_wholecell_mk_v14-ibiosim_dt0.1_tend50.0.csv']
[['time', '[A]', '[C]', '[D]'], ['time', 'A', 'C', 'D']]

# Elements
(501, 4)

# Maximum column difference
time    1.421085e-14
[A]     8.488962e-09
[C]     2.533890e-09
[D]     7.456576e-09
dtype: float64

# Maximum element difference
8.48896153371e-09

# Datasets are equal (diff <= eps=1e-05)
True
********************************************************************************
```

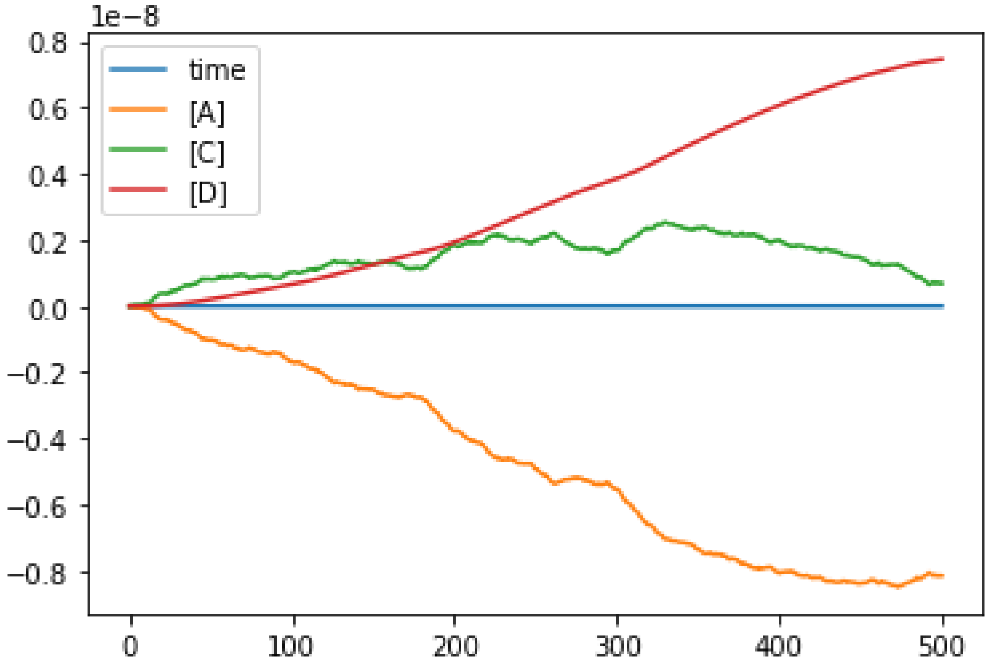

```
*********************************************************************************
['./toy_wholecell/toy_wholecell_mk_v14-sbmlutils_dt1.0_tend50.0.csv',
 './toy_wholecell/toy_wholecell_mk_v14-ibiosim_dt1.0_tend50.0.csv']
[['time', '[A]', '[C]', '[D]'], ['time', 'A', 'C', 'D']]

# Elements
(51, 4)

# Maximum column difference
time    0.000000e+00
[A]     1.280189e-07
[C]     5.191503e-08
[D]     1.242827e-07
dtype: float64

# Maximum element difference
1.28018868395e-07

# Datasets are equal (diff <= eps=1e-05)
True
*********************************************************************************
```

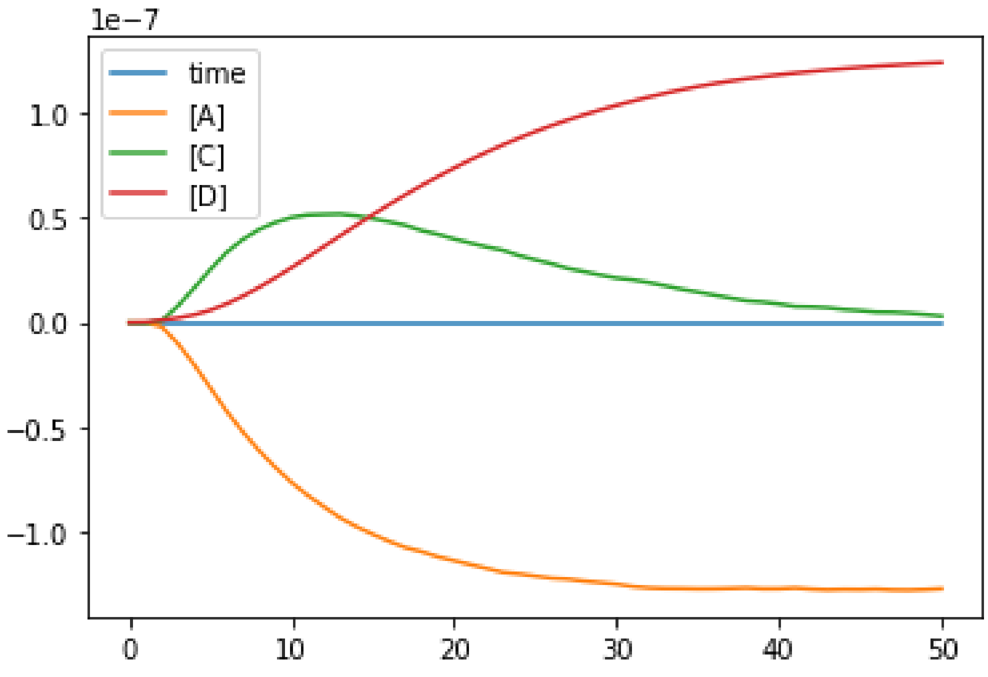

```
********************************************************************************
['./toy_wholecell/toy_wholecell_mk_v14-sbmlutils_dt5.0_tend50.0.csv',
 './toy_wholecell/toy_wholecell_mk_v14-ibiosim_dt5.0_tend50.0.csv']
[['time', '[A]', '[C]', '[D]'], ['time', 'A', 'C', 'D']]

# Elements
(11, 4)

# Maximum column difference
time    0.000000
[A]     0.000007
[C]     0.000003
[D]     0.000007
dtype: float64

# Maximum element difference
6.95619353408e-06

# Datasets are equal (diff <= eps=1e-05)
True
********************************************************************************
```

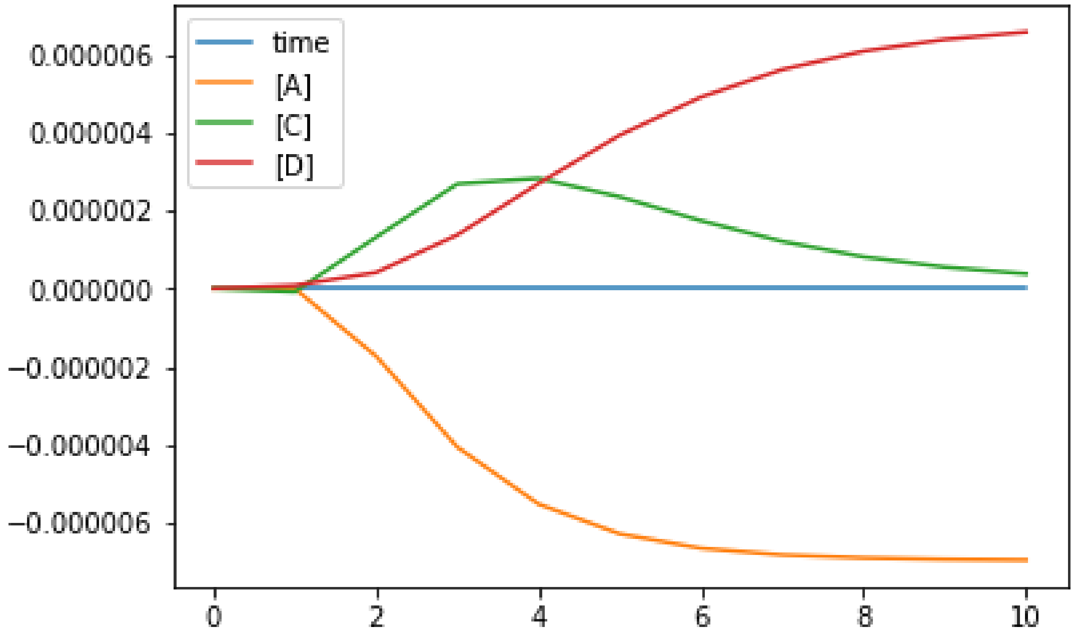

#### 1.2 toy_atp

```
In [3]: files = [
            "./toy_atp/toy_atp_mk_v12-sbmlutils_dt0.1_tend15.csv",
            "./toy_atp/toy_atp_mk_v12-ibiosim_dt0.1_tend15.csv"
        ]
        atp_dsc = DataSetsComparison(
            files=files,
            dfs = [
                pd.read_csv(files[0], sep="\t"),
                pd.read_csv(files[1])
            ],
            columns = [
                ["time", "[adp]", "[atp]", "[pyr]", "[glc]"],
                ["time", "adp", "atp", "pyr", "glc"],
            ]
        )

        atp_dsc.report()
```

```
********************************************************************************
['./toy_atp/toy_atp_mk_v12-sbmlutils_dt0.1_tend15.csv',
 './toy_atp/toy_atp_mk_v12-ibiosim_dt0.1_tend15.csv']
[['time', '[adp]', '[atp]', '[pyr]', '[glc]'],
 ['time', 'adp', 'atp', 'pyr', 'glc']]

# Elements
(151, 5)

# Maximum column difference
time     1.776357e-15
[adp]    3.505205e-07
[atp]    3.505205e-07
[pyr]    4.263256e-14
[glc]    1.781908e-14
dtype: float64

# Maximum element difference
3.50520538227e-07

# Datasets are equal (diff <= eps=1e-05)
True
********************************************************************************
```

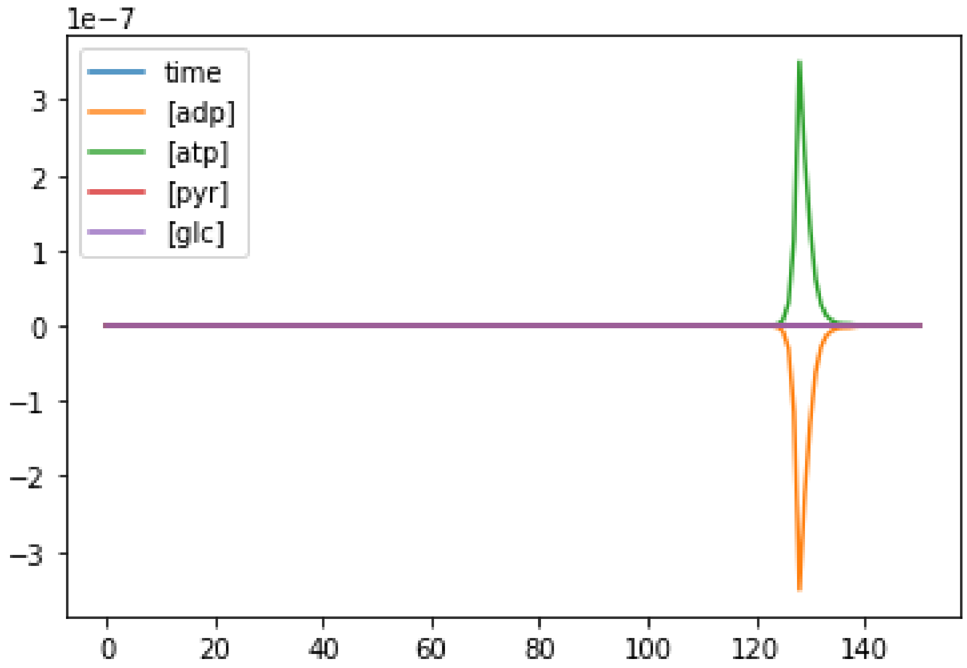


##### 1.3 diauxic

```
In [4]: files = [
            "./diauxic_growth/diauxic_growth_lw_v4-sbmlutils_dt0.01_tend15.0.csv",
            "./diauxic_growth/diauxic_growth_lw_v4-ibiosim_dt0.01_tend15.0.csv",
        ]
        diauxic_dsc = DataSetsComparison(
            files=files,
            dfs = [
                pd.read_csv(files[0]),
                pd.read_csv(files[1])
            ],
            columns = [
                ["time", "[Ac]", "[Glcxt]", "[O2]", "[X]"],
                ["time", "Ac", "Glcxt", "O2", "X"],
            ]
        )

        diauxic_dsc.report()
```

```
********************************************************************************
['./diauxic_growth/diauxic_growth_lw_v4-sbmlutils_dt0.01_tend15.0.csv',
 './diauxic_growth/diauxic_growth_lw_v4-ibiosim_dt0.01_tend15.0.csv']
[['time', '[Ac]', '[Glcxt]', '[O2]', '[X]'], ['time', 'Ac', 'Glcxt', 'O2', 'X']]
```

```
# Elements
(1501, 5)

# Maximum column difference
time        3.552714e-15
[Ac]        2.245718e-09
[Glcxt]     4.743860e-09
[O2]        1.326935e-09
[X]         3.114851e-10
dtype: float64

# Maximum element difference
4.74386013805e-09

# Datasets are equal (diff <= eps=1e-05)
True
********************************************************************************
```

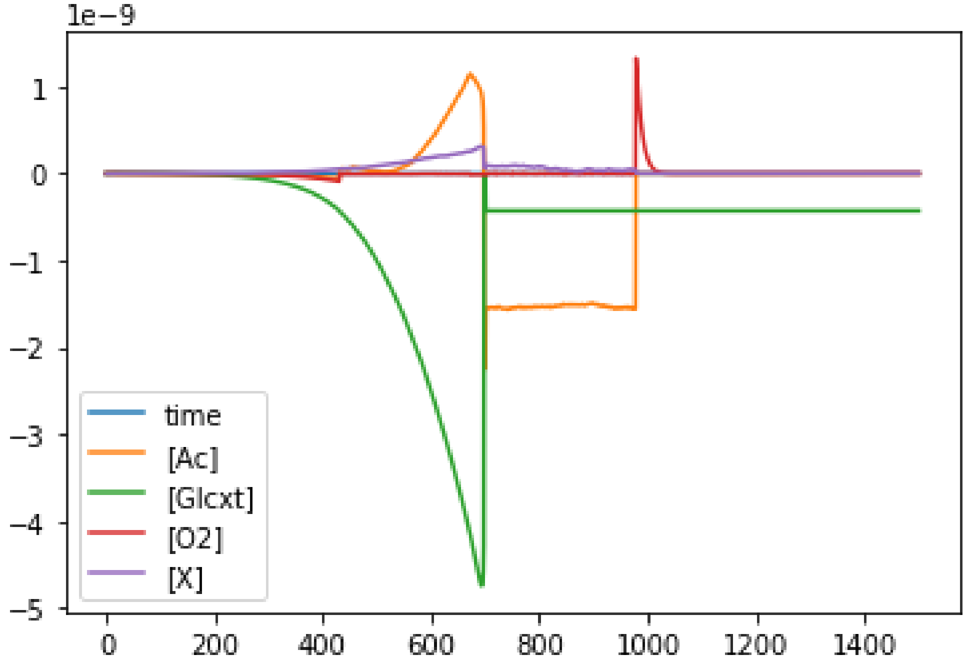


In [ ]:
